# Genetic Code Expansion and Enzymatic Modifications as Accessible Methods for Studying Site-Specific Post-Translational Modifications of Alpha-Synuclein and Tau

**DOI:** 10.1101/2025.05.16.654576

**Authors:** Ibrahim G. Saleh, Marie Shimogawa, Jennifer Ramirez, Bernard Abakah, Yarra Venkatesh, Honey Priya James, Ming-Hao Li, Sarah A. Louie, Marshall G. Lougee, Wai-Kit Chia, Richard B. Cooley, Ryan A. Mehl, Tobias Baumgart, Robert H. Mach, David Eliezer, Elizabeth Rhoades, E. James Petersson

**Affiliations:** Department of Chemistry, School of Arts and Sciences, University of Pennsylvania, 231 South 34th Street, Philadelphia, PA 19104, USA; Graduate Group in Biochemistry, Biophysics, and Chemical Biology, Perelman School of Medicine, University of Pennsylvania, 206 Anatomy-Chemistry Building, 3620 Hamilton Walk, Philadelphia, PA 19104, USA; Department of Biochemistry, Weill Cornell Medicine, 1300 York Avenue, New York, NY, 10065, USA; Department of Biochemistry and Biophysics Oregon State University, 2011 Ag Life Sciences Building, Corvallis, Oregon 97331, USA; Department of Radiology, Perelman School of Medicine, University of Pennsylvania, 231 S. 34th Street, Philadelphia, PA 19104, USA; Department of Biochemistry and Biophysics, Perelman School of Medicine, University of Pennsylvania, 421 Curie Boulevard, Philadelphia, PA 19104, USA

**Keywords:** Alpha-synuclein, tau, post-translational modification, genetic code expansion

## Abstract

Alpha-synuclein (αS) and tau play important roles in the pathology of Parkinson’s Disease and Alzheimer’s Disease, respectively, as well as numerous other neurodegenerative diseases. Both proteins are classified as intrinsically disordered proteins (IDPs), as they have no stable structure that underlies their function in healthy tissue, and both proteins are prone to aggregation in disease states. There is substantial interest in understanding the roles that post-translational modifications (PTMs) play in regulating the structural dynamics and function of αS and tau monomers, as well as their propensity to aggregate. While there have been many valuable insights into site-specific effects of PTMs garnered through chemical synthesis and semi-synthesis, these techniques are often outside of the expertise of biochemistry and biophysics laboratories wishing to study αS and tau. Therefore, we have assembled a primer on genetic code expansion and enzymatic modification approaches to installing PTMs into αS and tau site-specifically, including isotopic labeling for NMR and fluorescent labeling for biophysics and microscopy experiments. These methods should be enabling for those wishing to study authentic PTMs in αS or tau as well as the broader field of IDPs and aggregating proteins.

## Introduction

Alpha-synuclein (αS) and tau are classified as intrinsically disordered proteins (IDPs) that play crucial roles in cellular functions and are central to the pathology of numerous neurodegenerative diseases, particularly through their aggregation (**Figure 1**). In healthy contexts, αS exists primarily in the presynaptic terminals of neurons and its native roles are thought to be in vesicle trafficking and regulating neurotransmission (10; 9); 9). In pathological contexts, its fibrillar aggregates are in post-mortem brain tissue of patients with diseases collectively referred to as synucleinopathies, including Parkinson’s Disease (PD) and multiple system atrophy (MSA). Tau is a microtubule-associated protein found primarily in the axons of neurons thought to be involved in microtubule stabilization and axonal transport (31). Tauopathies, of which Alzheimer’s Disease (AD) is the most common, feature neuronal tau fibrils (33). There is also significant interest in non-fibrillar aggregates of αS and tau, including soluble oligomers and condensates (**Figure 1**). Critical to the functionality and dysregulation of IDPs are post-translational modifications (PTMs), particularly phosphorylation of Ser, Thr, or Tyr and Lys acetylation (**Figure 1**), which can influence interactions with other biomolecules and formation of pathological assemblies. Understanding the precise impact of site-specific PTMs on IDPs such as αS and tau is crucial for elucidating their roles in health and disease.

**Figure 1.**
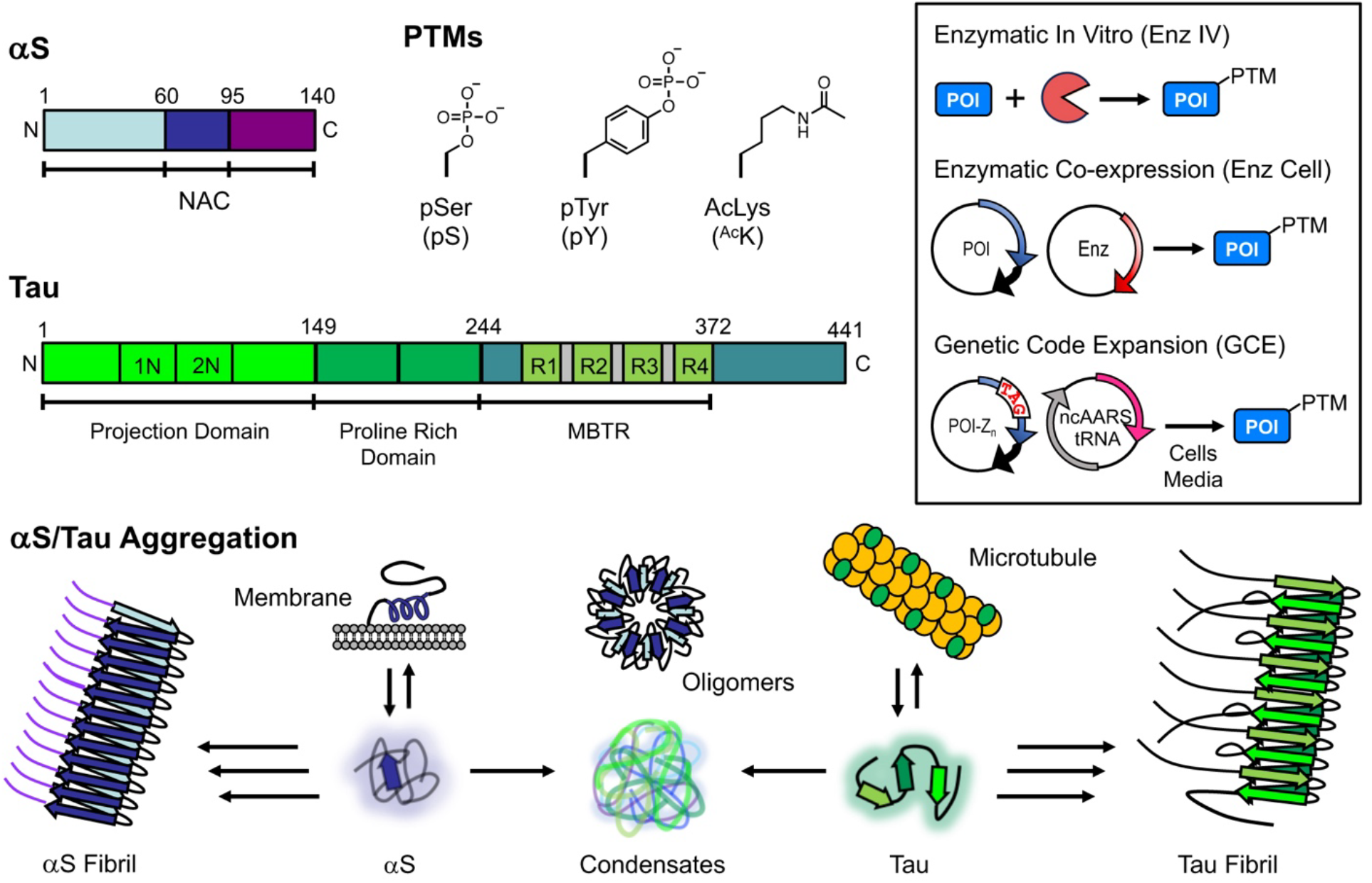
αS, Tau, PTMs, and Modification Methods. Top Left: αS and tau structures shown schematically with domains highlighted. PTM structures and abbreviations. Top Right (Inset): PTM installation methods for protein of interest (POI). Bottom: αS and tau functional states and aggregation pathways.

A large number of PTMs have been observed on αS and shown to affect its functional interactions with vesicles and pathological aggregation (41; 43; 25). Of these modifications, phosphorylation is one of the most prominent. Ser129 phosphorylation (pS_129_) is a hallmark of pathology, found in 90% of aggregated αS but only 4% of its soluble form, and is thought to disrupt dopamine uptake and inhibit αS binding to synaptic vesicles (24). While antibodies directed to pS_129_ are the standard in the field for observing PD and MSA pathology in immunohistochemistry, there are still questions as to its role in affecting αS function and aggregation. Tyr39 phosphorylation (pY_39_) has been associated with conflicting reports showing both increased neurodegeneration and slowed fibrillization kinetics in different studies (34; 6; 14). Inhibitors targeting the kinase responsible for Tyr39 phosphorylation, c-Abl, have undergone human trails as PD therapeutics, motivating a need for more a deeper understanding of pY_39_ effects on αS (5). Ser87 phosphorylation (pS_87_) has been shown to prevent αS fibril formation by increasing its flexibility as a monomer (38). Cryo-electron microscopy (cryo-EM) structures of αS fibrils composed of αS-pY_39_ or αS-pS_87_ have been solved, showing dramatic conformational changes relative to structures of unmodified fibrils (52; 27). However, these structures were determined with 100% modified protein, while studies of phosphorylation levels in patient tissue have shown that they are only present on 10-25% of αS (51). Thus, in spite of these studies clearly showing the impact of PTMs on αS function and pathology, there remain many unanswered questions needing further investigation.

PTMs have also been shown to play roles in the protein levels and aggregation of tau (42). Moreover, various PTMs have been detected in tau from healthy brains, suggesting their involvement in tau’s normal microtubule-associated functions (22). Indeed, about 35% of tau’s amino acid residues are susceptible to modification (2). One particular PTM of interest for tau is lysine acetylation, where recent reports have pointed to sex-specific impacts of Lys PTMs on tau. Despite women having higher tau levels and an increased risk of AD compared to men (12), the exact cause of this heightened susceptibility is not yet understood. It has been hypothesized that tau levels are regulated by an X-chromosome linked deubiquitinase that controls ubiquitin-mediated degradation through modification of Lys 274 and 281 (50). Tau acetylation should also play a regulatory role, since ubiquitination at these residues occurs in competition with lysine acetylation (^Ac^K). Additionally, ^Ac^K_274_, ^Ac^K_280_, and ^Ac^K_281_ are associated with impaired microtubule binding and increased tau aggregation (35; 11; 19; 22). Mass spectrometry (MS) identified Lys280 as a major site of tau acetylation in diseased tissue, suggesting its role in pathological tau transformation (11). Collectively, these studies prompt investigations of these three ^Ac^K sites to determine how much of the impact of acetylation comes from direct functional modulation versus its ability to block ubiquitination.

Traditionally, biochemists and biophysicists have studied PTMs through a strategy of mutating the target sites to other naturally occurring amino acids that mimic the desired PTM. Commonly used “phosphomimetics” include glutamate or aspartate; however, they lack the dianionic nature of the phosphate group (49) and are less sterically bulky than tyrosine, which can result in different properties compared to phosphotyrosine (40). To investigate acetylation, lysine is often mutated to glutamine, but this involves a significant reduction in sidechain length. These mimics, while easy to introduce, often do not reproduce the effects of the authentic PTM, particularly in IDPs like αS and tau, where single point mutations can produce dramatic changes in folding and dynamics (41).

The study of authentic, site-specific PTMs in proteins has largely been approached through various methods including site-specific mutagenesis, chemical or semi-synthesis, and enzymatic reactions, each with inherent limitations. While mutagenesis allows for facile introduction of mutation-based mimics of PTMs, these alterations do not always recapitulate the chemical and structural nuances of naturally occurring PTMs (40). On the other hand, the use of peptide ligation techniques, although precise, often require sophisticated synthetic expertise and can be resource and time intensive (36). Enzymatic methods are limited by the availability of specific, efficient enzymes, but have proven successful in certain instances where site-specificity and yield satisfy the experimental needs (28; 40; 44). It must be noted that enzymatic modification often requires additional mutations to eliminate off-target introduction of PTMs, such as Ser-to-Ala or Tyr-to-Phe mutations to prevent phosphorylation, and these mutations can introduce artifacts (3). We highlight cases involving αS where enzymatic methods have effectively introduced PTMs, either through in vitro enzymatic treatment (Enz IV) of the purified protein or through co-expression of the enzyme in cells (Enz Cell) by co-transformation of a plasmid for the protein of interest (POI) and a plasmid for the enzyme (**Figure 2**).

**Figure 2.**
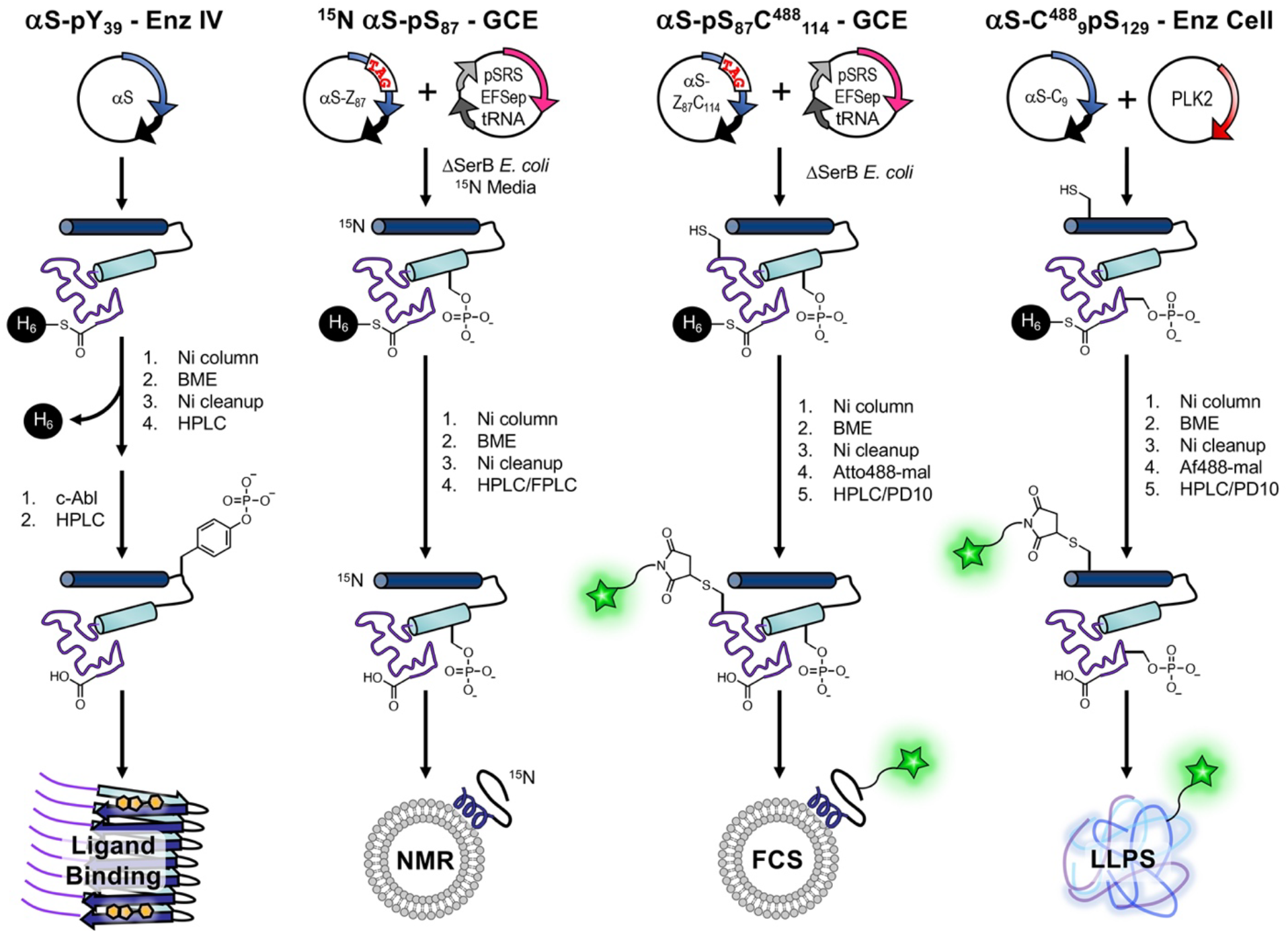
Introduction of PTMs and Labels to αS for Diverse Applications. Enz IV, GCE, and Enz Cell workflows shown, including labeling ^15^N labeling for NMR and fluorescent labeling for FCS or LLPS microscopy. Intein six His (H_6_) tags are used for initial purification followed by more intensive HPLC/FPLC purification or simpler PD10 column purification for dye removal.

For broader applicability and ease of use, genetic code expansion (GCE) is a robust alternative (13). GCE involves the targeted incorporation of noncanonical amino acids (ncAAs) into proteins, typically by mutation to introduce a TAG (amber) stop codon at the site of interest. Delivery of the ncAA is achieved through an orthogonal tRNA_CUA_, which recognizes the TAG codon, and an engineered orthogonal aminoacyl-tRNA synthetase (RS) designed or evolved for the specific ncAA of interest (15). GCE incorporation of the PTM is then performed by co-transformation of a plasmid for the POI with a TAG codon at the PTM site and a plasmid for the ncAARS and tRNA_CUA_, sometimes in combination with specialized cells or media (**Figure 2**). Cooley and Mehl have developed an efficient phosphoserine GCE system (pSer-3.1G) using a robust release factor 1 deficient *E. coli* strain to minimize truncation issues that can arise from reassignment of the TAG stop codon (54; 1). For ^Ac^K incorporation, we use an RS system developed by Liu and coworkers using phage-assisted continuous evolution (16; 8), chAcK3RS with IPYE mutations, based on the *M. barkerii* pyrolysyl RS (37). Although neither system had been previously used with αS and tau, it was relatively easy to adapt them for use with these proteins, highlighting the versatility of the GCE approach.

Herein, we showcase the use of either enzymatic modifications or GCE to study four PTM targets: i) pS_87_ in αS (GCE); ii) pY_39_ in αS (Enz IV); iii) pS_129_ in αS (GCE and Enz Cell); and iv) ^Ac^K_274_, ^Ac^K_280_, and ^Ac^K_281_ in tau (GCE). We then characterize how these PTMs affect factors critical in αS and tau native or pathological contexts using exemplary assays: αS vesicle binding through fluorescence correlation spectroscopy (FCS) and NMR, αS fibril conformation through radioligand binding, tau-mediated tubulin polymerization, and phase transition through fluorescence microscopy of mixed αS/tau condensates formed by liquid/liquid phase separation (LLPS). By integrating successful enzymatic modifications with GCE, we aim to provide guidance for biochemists and biophysicists who wish to achieve a nuanced understanding of authentic PTM impacts on IDPs.

### Production of phosphorylated αS by GCE

Using methods similar to those that we have previously described for recombinantly expressing αS with ncAAs in *E. coli*, one can produce αS with phosphorylation of serine 87 (αS -pS_87_) as shown in **Figure 2**. We use two plasmids, one encoding the aaRS for phosphoserine and its cognate tRNA as well as EFSep, an EF-Tu mutant that better accommodates delivery of pSer-aminoacylated tRNA, the other encoding the POI with a TAG stop codon at the site of ncAA incorporation. Often, our POI bears a C-terminal traceless intein-His_6_ tag which not only allows for isolation of full-length products from species truncated at the TAG codon but could also be cleaved off with thiols during purification and provide scarless protein products (4).

For pSer GCE, *E. coli* cells with a ΔserB phosphatase genomic knockout (BL21 ΔserB or B95 ΔserB cells developed by Cooley et al. (54), the latter lacks release factor 1) were used so that pSer is accumulated intracellularly and readily available for GCE, since pSer cannot be exogenously added due to its low cell permeability. Following affinity purification, high performance liquid chromatography (HPLC) was used to separate the product from impurities, which were characterized by matrix-assisted laser desorption ionization mass spectrometry (MALDI-MS) and found to include de-phosphorylation (−80 Da), Gln misincorporation, and an unknown +72 Da species (**Figure 3** and **Figure S3**). We found BL21 ΔserB cells to be higher yielding than B95 ΔserB cells, with a 2-fold higher level of protein expression and slightly lower fraction of impurities. As a method to determine the homogeneity of phosphorylated samples, we performed Phos-tag gel electrophoresis, where the phosphate-binding ligand usually slows the mobility of phosphorylated proteins (32). We observed a mobility shift of αS-pS_87_, where the extent of the shift seems to depend slightly on buffer components (**Figure S1**, see marked bands in “Ni cleanup” and “HPLC purified”). We observed nearly quantitative isolation of phosphorylated product from the unphosphorylated species by HPLC (**Figure 3** and **Figure S2**, HPLC, **Figure S3**: MALDI-MS of HPLC-purified product).

**Figure 3.**
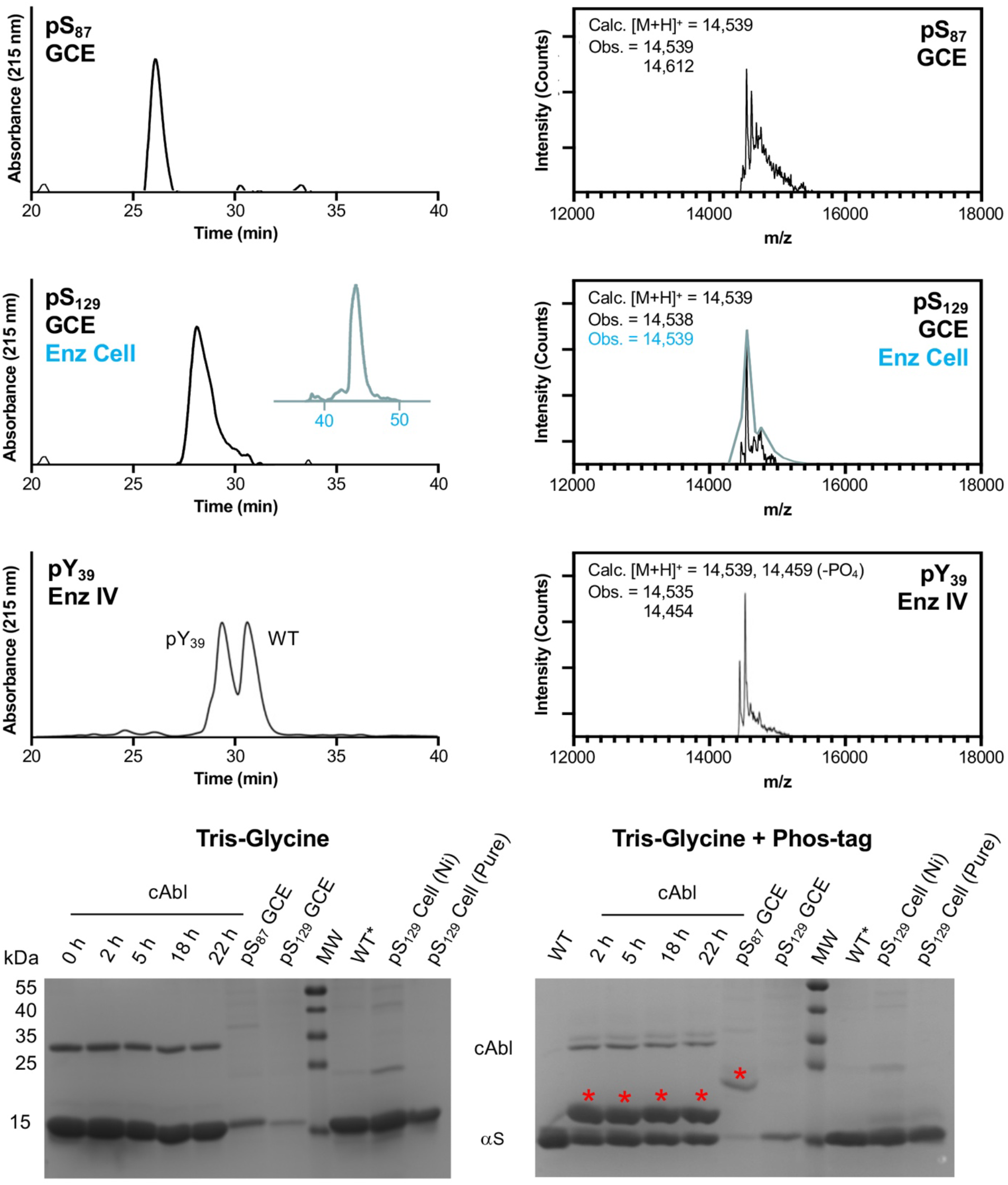
Characterization of Phosphorylated αS. Top: HPLC and MALDI MS. Bottom: Phos-tag gel analysis of αS constructs after treatment of purified WT with c-Abl for varying amounts of time, after purification from GCE expression, or after purification from enzymatic co-expression in cells. αS-pS_129_ from cellular expression (teal) was analyzed using a different HPLC gradient than αS-pS_129_ from GCE (black). WT* indicates N-terminally acetylated αS, produced by enzymatic expression as previously described. αS-pS_129_ produced by enzymatic co-expression was tested after Ni column elution, and also after additional HPLC purification. Red asterisks indicate bands for phosphorylated αS on the Phos-tag gel.

We expressed αS-pS_129_ using the pSer-3.1G system as well, and similarly observed a mixture of products, with the predominant species being the desired phosphorylated protein (**Figure 3**). However, unlike αS-pS_87_, the αS-pS_129_ species could not easily be separated by HPLC and yields (10.2 mg/L) were limited by the need to collect fractions judiciously to obtain pure phosphorylated protein. Interestingly, we did not observe significant amounts of the +72 Da species for incorporation at Ser129. In spite of MALDI-MS data indicating the presence of pSer, we observed no shift on the Phos-tag gel for αS-pS_129_ (**Figure 3**), a finding corroborated by production of αS-pS_129_ through co-expression with a kinase (see below). This is likely due to the acidic nature of the C-terminal region of αS.

To demonstrate the power of the GCE method over NCL-based incorporation, we additionally prepared a fluorescently labeled αS-pS_87_ construct for FCS and an isotopically labeled of αS-pS_87_ construct for NMR. For FCS, fluorescently labeled αS variants with a Cys mutation at site 114 were prepared by site directed mutagenesis, and reacted with Atto488-maleimide overnight at room temperature, resulting in labeled, phosphorylated construct (αS-pS_87_C^488^_114_) in 0.5 mg/L yield. This illustrates the value of the GCE approach, as we had previously synthesized αS-pS_87_C^488^_114_ by NCL, but obtained yields too low for full characterization of vesicle binding curve (18). For NMR, we prepared ^15^N isotopically labeled αS-pS_87_, a construct that was prohibitively expensive to produce by NCL due to the high cost of isotopically labeled amino acids for solid phase peptide synthesis. The ^15^N αS-pS_87_ expression followed strategies introduced by Cooley et al. (48) – it was necessary to start the culture with rich media and then switch to a minimal media so that Ser auxotroph *E. coli* ΔserB cells were kept alive while minimizing ^14^N incorporation. The proteins were expressed at lower temperature, but for longer, to minimize the activity of endogenous phosphatases. HPLC was again used to purify phosphorylated αS (0.3 mg/L yield) from de-phosphorylated side products, wherein we observed that a larger fraction of αS was de-phosphorylated in the ^15^N isotopic labeling protocol (**Figure S2**: HPLC, **Figure S3**: MALDI-MS of HPLC-purified product).

### Production of phosphorylated αS by enzymatic modification

For comparison to GCE, production of αS-pS_129_ was also performed as in previous reports from the Rhoades laboratory (44), by co-expressing αS with Ser/Thr kinase PLK2 in *E. coli*. This is fortunately possible thanks to the ability of PLK2 to phosphorylate Ser129 selectively and quantitatively without posing toxicity to *E. coli*. This method gives a very high yield (6.9 mg/L yield) and therefore for this phosphorylation site, this was our method of choice over the GCE approach. Again, we observed no mobility shift of this phosphorylated product on Phos-tag gel, when tested either immediately after Ni column elution (to minimize any potential phosphate hydrolysis or elimination) or after further purification (**Figure 3**). To generate fluorescently labeled αS-pS_129_ for microscopy studies, we expressed αS-C_9_pS_129_ using the PLK co-expression method (5.9 mg/L yield), purified the protein using a Ni column, and reacted it with Alexafluor 488-maleimide, followed by HPLC purification to generate αS-C^488^ _9_pS_129_ (2.6 mg/L yield), which was characterized by MALDI-MS (**Figure S6**).

Production of αS-pY_39_ was performed using an *in vitro*, chemoenzymatic approach with a recombinantly expressed catalytic domain of the tyrosine kinase c-Abl, similar to our published approach. It has been noted that in prokaryotes like *E. coli*, c-Abl needs to be co-expressed with the phosphatase YopH to counteract and suppress its toxicity, and this makes it impossible to co-express the kinase with αS and get Tyr39 phosphorylated at appreciable levels in *E. coli* cells (39). In previous reports, we used a semi-synthetic approach to ensure that αS was phosphorylated only at Tyr39. Here, in order to avoid the need for protein ligation we performed enzymatic modification on full-length αS. We chose to do this because we were able to achieve an acceptable level of conversion and site-selectivity of c-Abl for Tyr39 phosphorylation. Upon incubation of αS with c-Abl, we observed a mobility shift for ∼50% of αS on Phos-tag gel, with confirmatory analysis by HPLC and MALDI-MS (**Figure 3**). Following Glu-C digestion and MALDI-MS analysis, we observed only the αS_36-46_ peptide as a phosphorylated peptide, where MS/MS analysis showed that Tyr39 was phosphorylated (**Figure S5**). As one can see in the HPLC trace in **Figure 3**, αS-pY_39_ can be separated from unmodified αS if one wishes to analyze 100% phosphorylated material (Note: Separation can be performed by analytical HPLC, but obtaining pure αS-pY_39_ may not be possible at the semi-prep scale). In our case, we were interested in more physiologically-relevant levels of phosphorylation (10-25% at Tyr39, as determined from analyses of patient samples) (51), so we simply mixed the 50% modified protein with unmodified αS prior to aggregation to form fibrils.

### Production of acetylated tau by GCE

To produce acetylated tau constructs, plasmids encoding the 0N4R tau variant with a TAG mutation at Lys274, Lys280, or Lys281 were generated by site-directed mutagenesis. Each tau TAG mutant plasmid was co-transformed into BL21 *E. coli* cells with the plasmid encoding the chAcK3RS/tRNA_CUA_ pair and expressed in media with ^Ac^K and 50 mM nicotinamide (to suppress deacetylases). The tau plasmid used in these experiments has an N-terminal His tag with a tobacco etch virus (TEV) cleavage site. After Ni column purification, the His tag was cleaved with TEV and the tau AcK constructs were purified by fast protein liquid chromatography (FPLC) using a size exclusion column to obtain tau-^Ac^K_274_, tau-^Ac^K_280_, and tau-^Ac^K_281_ in 1.1, 1.2, and 1.2 mg/L yields, respectively. Acetylation was confirmed via MALDI-MS and ESI-MS to be ≥95% for all three constructs (**Figure 4 and Figure S7**).

**Figure 4.**
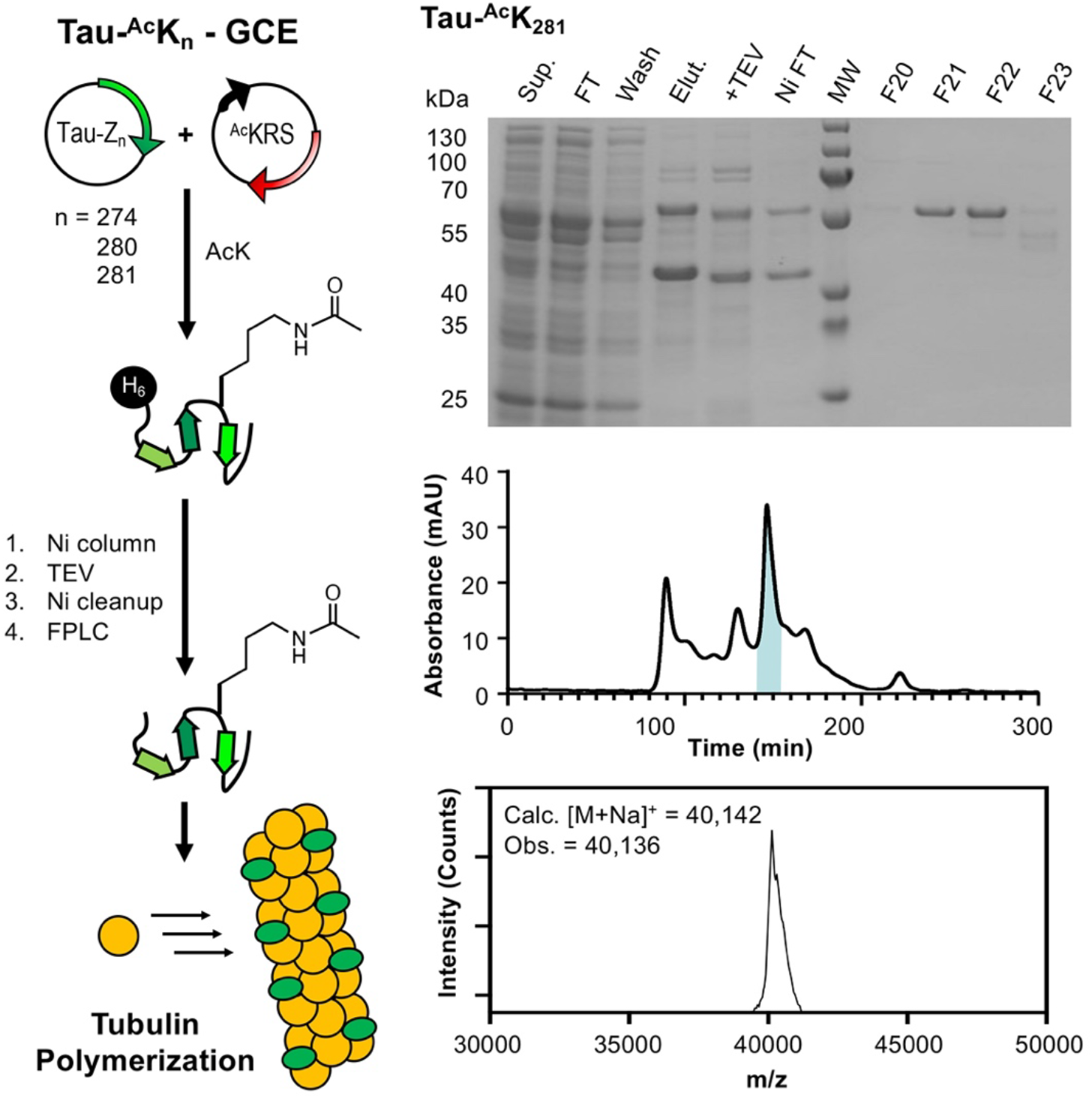
Expression and analysis of tau-^Ac^K_281_ construct. Left: Expression of acetylated tau constructs and use in tubulin polymerization assays. Top Right: Gel analysis of expression and purification. Middle Right: FPLC purification. Highlighted fractions correspond to F20-F23 shown on gel above. Bottom Right: MALDI MS analysis of F21 and F22 confirming the identity of tau-^Ac^K_281_.

With these purified, PTM-modified αS and tau constructs, we then measured αS-pS_87_ vesicle binding through FCS and NMR, studied αS-pY_39_ fibril conformation through radioligand binding, probed tau acetylation impacts on tubulin polymerization, and imaged the effects of Ser129 phosphorylation on αS/tau condensate formation.

### Effects of pY_39_ on radioligand binding to αS fibrils

Radioligands, such as positron emission tomography (PET) tracers, are powerful tools for imaging αS aggregates in tissues and *in vivo* that can provide insights into misfolding and aggregation in PD and MSA. The Petersson and Mach laboratories have developed PET probes to bind selectively and with high affinity to specific sites on αS fibrils in order to visualize disease-associated aggregates (26; 17). Binding sites were computationally and experimentally mapped based on the original solid-state NMR structure (PDB ID: 2n0a) published by Rienstra and coworkers, which is similar to many unmodified αS fibril structures published to date, including ones solved by cryo-EM (41). Building on this, one could take radioligand PET probes with known binding sites in unmodified αS fibrils (Tg-190b for “Site 2” and BF-2846 for “Site 9”) and investigate their binding to post-translationally modified αS fibrils, then use this information to better understand the conformational differences introduced by specific modifications. Since PTMs can significantly alter αS fibril structure and behavior, radioligand binding provides a functional readout that can hint at how these modifications influence fibril conformation. αS-pY_39_ was of particular interest, not only because Tyr39 is within Site 2, the target of Tg-190b binding, but also because Zhao et al. showed through protein synthesis and cryo-EM that the structure of 100% αS-pY_39_ fibrils (PDB IDs: 6l1t and 6l1u) is dramatically different than unmodified, wild type (WT) fibrils (53). However, since physiological αS-pY_39_ levels range from 10 to 25% (51), we were interested in whether these lower levels of PTM modification induced a conformational change that could be detected by radioligand binding.

Fibrils were prepared from 100% WT αS or mixtures with 10% or 25% αS-pY_39_. Binding of Tg-190b, the Site 2 binder, was significantly disturbed by the presence of pY_39_ with the apparent dissociation constant (K_d_) increasing from 5.2 nM to 19 nM to 35 nM with increased phosphorylation (**Figure 5**), whereas the K_d_ BF-2846, the Site 9 binder, was about 1 nM for all three fibril samples. This suggests that at physiological stoichiometries, αS-pY_39_ fibrils have a similar structure to that of WT fibrils in the Site 9 region, with only local effects around Site 2. This finding is incompatible with the broad conformational rearrangement observed by Zhao et al. in 6l1t/6l1u or mixtures of the 2n0a and 6l1t/6l1u fibril polymorphs, both of which would dramatically change Site 9 binding since the pocket around Phe94 is eliminated in 6l1t/6l1u (**Figure 5**). Indeed, the 10-fold change in K_d_ for Site 2 also seems to be less dramatic than what one would expect for Tg-190b binding to the 6l1t/6l1u polymorph, since pY_39_ is buried in the fibril core, stabilized by salt bridge interactions with several Lys residues. While further investigations using ssNMR or cryo-EM will be necessary to fully understand the structural basis for these effects on ligand binding, these experiments demonstrate the value of accessing modified αS to investigate the impact of physiological PTMs on ligand binding.

**Figure 5.**
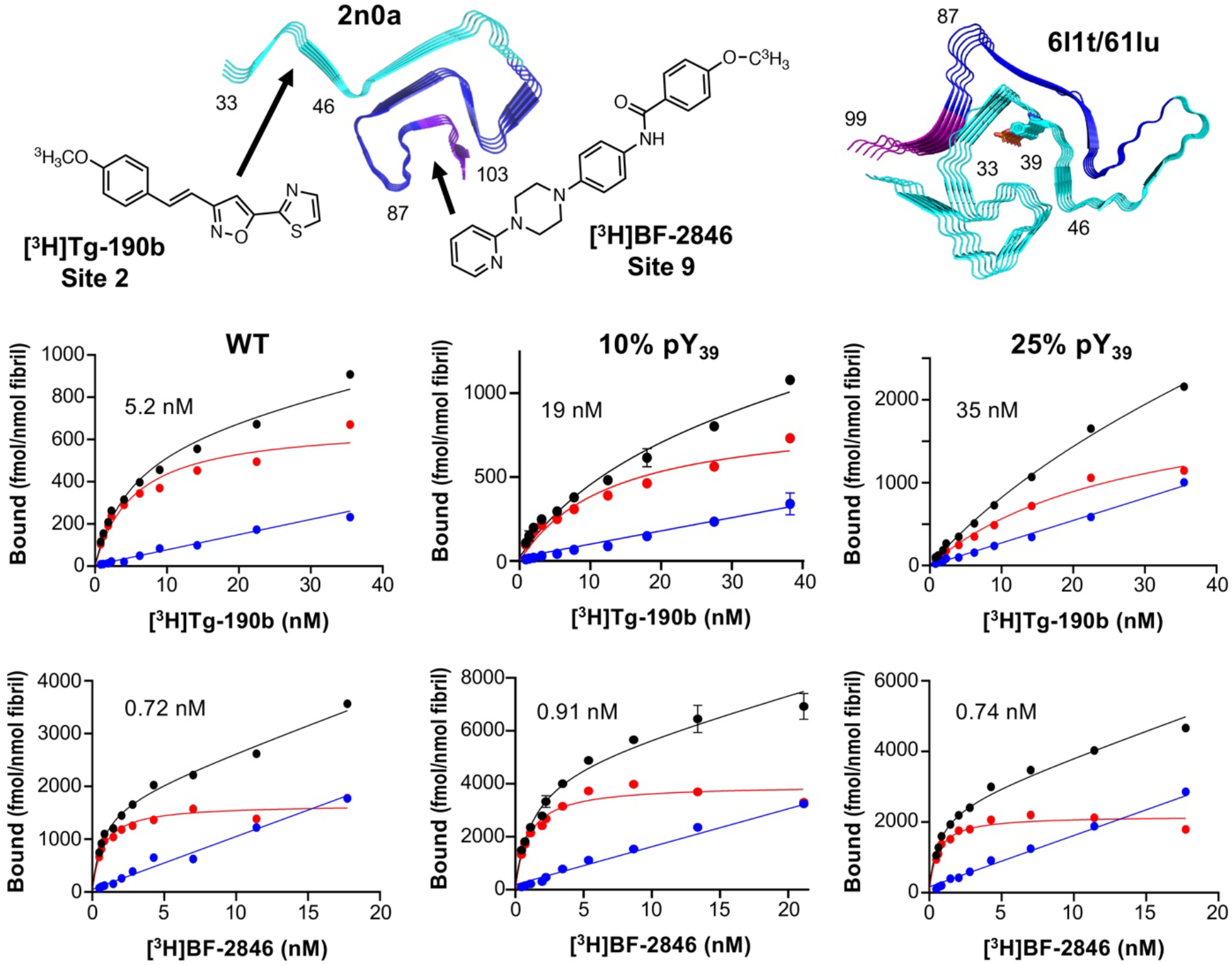
Radioligand binding to fibrils containing pY_39_ at physiological levels. Top: Site 2 (residues 33-46) and Site 9 (residues 87-103) shown on the ssNMR structure of WT αS fibrils (PDB ID: 2n0a) and the cryo-EM structure of 100% αS-pY_39_ fibrils with the structures of Site selective ligand Tg-190b and Site 9 selective ligand BF-2846. Bottom: Saturation binding of [^3^H]Tg-190b and [^3^H]BF-2846 radioligands to fibrils made with 100% WT αS, 10% αS-pY_39_, or 25% αS-pY_39_ shows that Site 2 is perturbed while Site 9 is not.

### Biophysical characterization of vesicle binding of αS-pS_87_

Phosphorylation at Ser87 has been studied by multiple groups, with a focus on aggregation -the Lashuel group showed that pS_87_ levels are increased in synucleinopathy patient brains, and performed *in vitro* and rat studies, where it slowed down and reduced aggregation, resulting in less toxicity (38). Notably, they used either mutational mimics or enzymatic phosphorylation by kinase CK1 coupled to a S_129_A mutation to block phosphorylation at that site. The Churchill Group used GCE and incorporated authentic pS_87_ modification with minimal sequence scars from TEV cleavage, which exhibited dopamine or Cu^2+^-induced oligomerization trends that were similar to WT, but significantly increased toxicity in SH-SY5Y neuronal cell model (21). Lastly, the Li and Liu groups performed protein semi-synthesis, resulting in a completely authentic αS-pS_87_ construct, and their a cryo-EM study showed its unique fibril structure – unlike the case of pY_39_, pS_87_ could not be stabilized by nearby positively charged residues and causes the broader C-terminal region of αS to be excluded from the fibril core while including the entire N-terminal region (PDB: 8jey) (27).

On the other hand, the effects of pS_87_ on the native role of αS are relatively understudied: the Lashuel group studied αS conformation in the presence and absence of lipid membranes using circular dichroism, indicating that pS_87_ alters the conformation of the helix that forms when bound to SDS micelles and that it reduces helicity on vesicles (38). Again, they used the proteins modified with mutational mimics/blocks and enzymatic modification for this.

To understand the effects of authentic pS_87_ on vesicle binding, we first acquired proton-nitrogen heteronuclear single quantum coherence spectra (^1^H,^15^N – HSQC) in the presence and absence of small, unilamellar vesicles (SUVs) that are composed of 60:25:15 1,2-dioleoyl-sn-glycero-3-phosphocholine/1,2-dioleoyl-sn-glycero-3-phosphatidylethanolamine/1,2-dioleoyl-sn-glycero-3-phospho-L-serine (DOPC/DOPE/DOPS). The NMR resonance chemical shift changes between spectra for free αS WT or αS-pS_87_ were very minimal (**Figure S9**). We observed that both pS_87_ and WT led to a ∼60% reduction of intensity for resonances corresponding to αS residues ∼1-100 (**Figure 6**). This reduction is caused by binding of this region to vesicles, which are slowly tumbling. The lack of any difference in the binding profiles suggests that pS_87_ does not substantially affect vesicle binding under these conditions.

**Figure 6.**
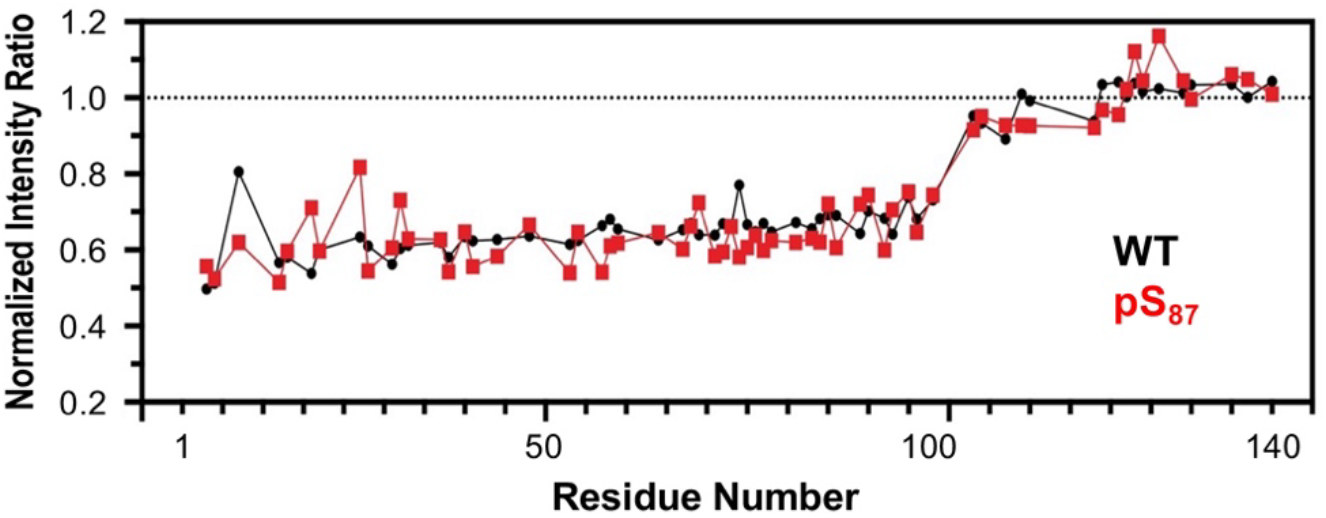
Effects of pS_87_ on vesicle binding, investigated by NMR. NMR intensity ratios for each residue were calculated from ^1^H-^15^N HSQC spectra collected with ^15^N-labeled αS variants in the presence or absence of SUVs, normalized by the average ratio for residues 101-140. No significant differences in binding are seen.

To rigorously determine the effect of pS_87_ on the αS vesicle K_d_, we performed FCS, which is a well-established method for quantifying biomolecule interactions, including αS vesicle binding (46). FCS and NMR are complementary techniques in this context: FCS provides quantitative measurements with a slight perturbation to the protein, while NMR offers qualitative insights and low-resolution structural information without perturbation. It is also notable that this binding affinity measurement could not be completed when we previously generated pS_87_ by chemical protein synthesis due to low yields (18). For this experiment, we used fluorescently labeled αS variants with a Cys mutation at site 114, αS-C^488^_114_ and αS-pS_87_C^488^_114_ (denoted WT and pS87, respectively, in **Figure 7**). We prepared lipid vesicles composed of 50:50 1-palmitoyl-2-oleoyl-sn-glycero-3-phospho-L-serine/1-palmitoyl-2-oleoyl-glycero-3-phosphocholine (POPS/POPC). The diffusion times of the free αS proteins and vesicles were initially measured. To evaluate αS-vesicle binding, a fixed amount of labeled αS construct was added to varying concentrations of vesicles, and the fraction of protein bound was determined by fitting a two-component autocorrelation function. The fraction bound at each vesicle concentration was then used to generate a binding curve for each αS construct. We observed no significant difference in vesicle binding affinity caused by phosphorylation at Ser87 (**Figure 7**; K_d,app_(WT) = 3.7 ± 0.5 μM, K_d,app_(pS_87_) = 5.2 ± 0.5 μM). This result is consistent with our observations in the NMR vesicle binding experiments, showing the complementarity of these methods, both using PTM-modified αS constructs that are difficult to access through NCL.

**Figure 7.**
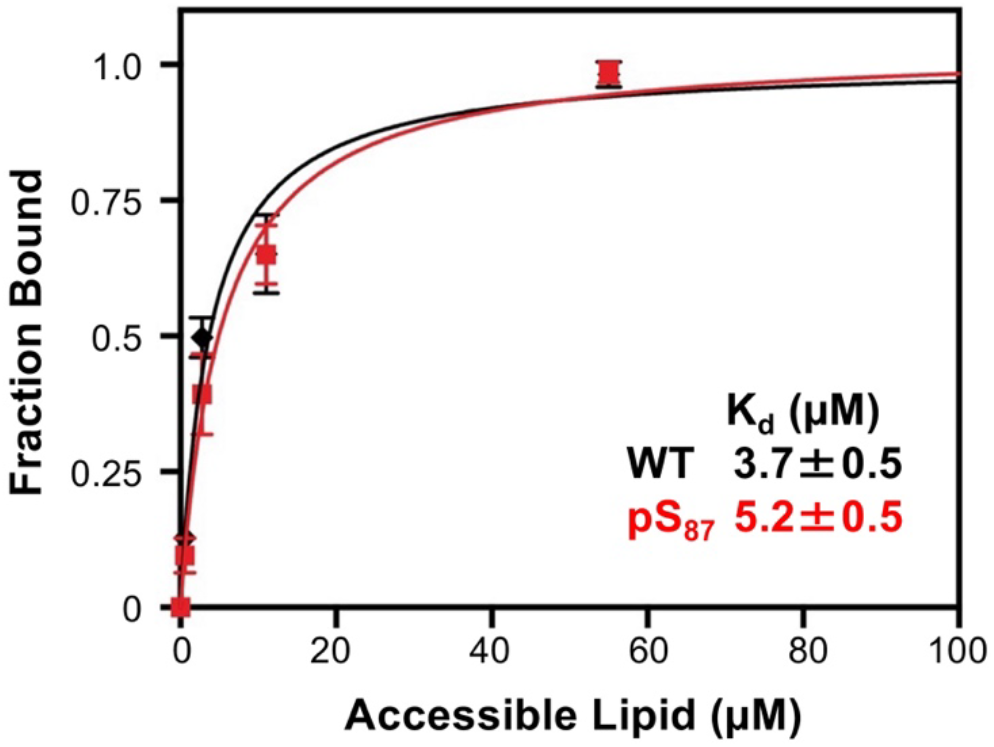
Effects of pS_87_ on vesicle binding, investigated by FCS. αS constructs labeled with Atto488 were examined in the presence of varying concentrations (1 to 50 µM accessible lipid) of lipid vesicles consisting of 50:50 POPS/POPC, showing no significant effect on the apparent K_d_.

### Tau acetylation effects on microtubule polymerization

K_274_, K_280_, and K_281_ have been identified as pathologically relevant sites through mass spectroscopy studies of diseased tissue. It is critical to investigate the potential functional and regulatory roles of acetylation in tau-microtubule interactions. Thus, microtubule polymerization assays were conducted on the three acetylated 0N4R tau constructs generated using GCE (^Ac^K_274_, ^Ac^K_280_, ^Ac^K_281_), as well as unmodified 0N4R tau (WT). These polymerizations were conducted at 37 °C, a constant GTP concentration, and a 1:2 concentration ratio of tau to tubulin. Light scatter at 340 nm was used to quantify the increasing turbidity of the solution as microtubule polymerization occurred. Spontaneous tubulin/GTP polymerization in the absence of tau was measured to serve as a control. The ^Ac^K_274_ variant induced polymerization at a similar rate to WT, but both ^Ac^K_280_ and ^Ac^K_281_ showed a slight reduction in the polymerization rate (**Figure 8**).

**Figure 8.**
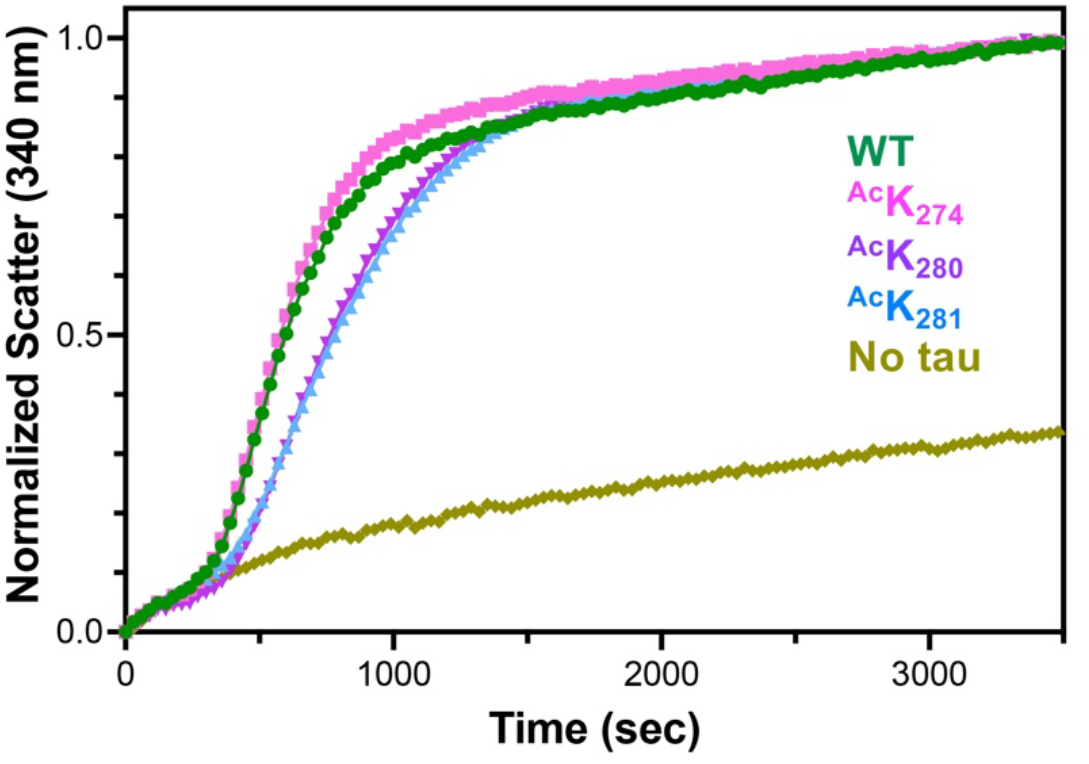
Microtubule polymerization assay with acetylated tau. WT 0N4R tau (green) showed comparable polymerization rates to tau-^Ac^K_274_ (pink). Tau-^Ac^K_280_ (teal) and tau-^Ac^K_281_ (blue) both showed slightly impaired polymerization compared to WT. Tubulin/GTP-only (mustard) was included as a negative control experiment. Polymerization data are an average of 3 replicates per day with 3 overall replicates.

Although each construct had only a single acetylation site, we were surprised to observe a greater impairment in polymerization rate for ^Ac^K_280_/^Ac^K_281_ compared to ^Ac^K_274_. These results suggest a connection to the resolved near-atomic cryo-EM model of the microtubule-tau complex by Kellogg et al. which demonstrated that K_281_ plays a crucial role in the complex (30). Further structural work by Brotzakis et al. quantified tau-microtubule contact interactions and found that K_274_ has much fewer contacts than K_280_ and K_281_, suggesting that acetylation of those two sites would have a larger impact on the microtubule-tau complex, as observed in the polymerization kinetics (7). Additionally, Yan et al. (50), demonstrated that in HEK293T cells co-transfected with tau K_280_Q or K_281_Q and USP11, there was reduced polyubiquitin, whereas K_274_Q cells showed no change. This suggests that the ^Ac^K_280_ and ^Ac^K_281_ sites might have a more pathological role, while ^Ac^K_274_ may act as a regulatory buffer depending on the need for acetylation or ubiquitination. Yan et al. also found that USP11 siRNA fully prevented the increase in ^Ac^K_281_ and partially suppressed the increase in ^Ac^K_274_ induced by inhibition of lysine deacetylase SIRT1, highlighting the distinct roles of the 280/281 sites versus the 274 site (50).

### Effect of pS129 on size distribution and dynamics of αS in αS/tau co-condensates *in vitro*

As a final example of biophysical studies enabled by production of PTM-modified proteins, we investigated αS/tau co-condensate formation. Recently, αS has been shown to interact with tau through electrostatic attractions, forming LLPS condensates between the highly negatively charged C-terminal region of αS and the positively charged proline-rich region of tau (20). To examine the impact of the pS_129_ modification on the size distribution and dynamics of αS in co-condensates with tau, we incubated αS-C^488^_9_ (WT) or αS-C^488^_9_pS_129_ (pS_129_) with BODIPY 558-labeled 0N4R tau (tau^BDP558^), with each labeled protein present in a 5:95 mixture with the corresponding unlabeled construct (**Figure 9**). Using two-color confocal microscopy, we confirmed the formation of αS/tau co-condensates, detecting αS in the 500–540 nm channel and tau in the 570–620 nm channel. The size distribution of WT/tau and pS_129_/tau co-condensates showed no significant difference, with average diameters of 1.33 ± 0.36 µm and 1.36 ± 0.34 µm, respectively. However, fluorescence recovery after photobleaching (FRAP) revealed that pS_129_ modification alters the dynamics of αS within αS/tau co-condensates. Specifically, pS_129_ αS exhibited a significantly slower fluorescence recovery time (τ_FR_ = 22.32 ± 8.73 s) compared to WT αS (τ_FR_ = 6.03 ± 1.67 s).

**Figure 9.**
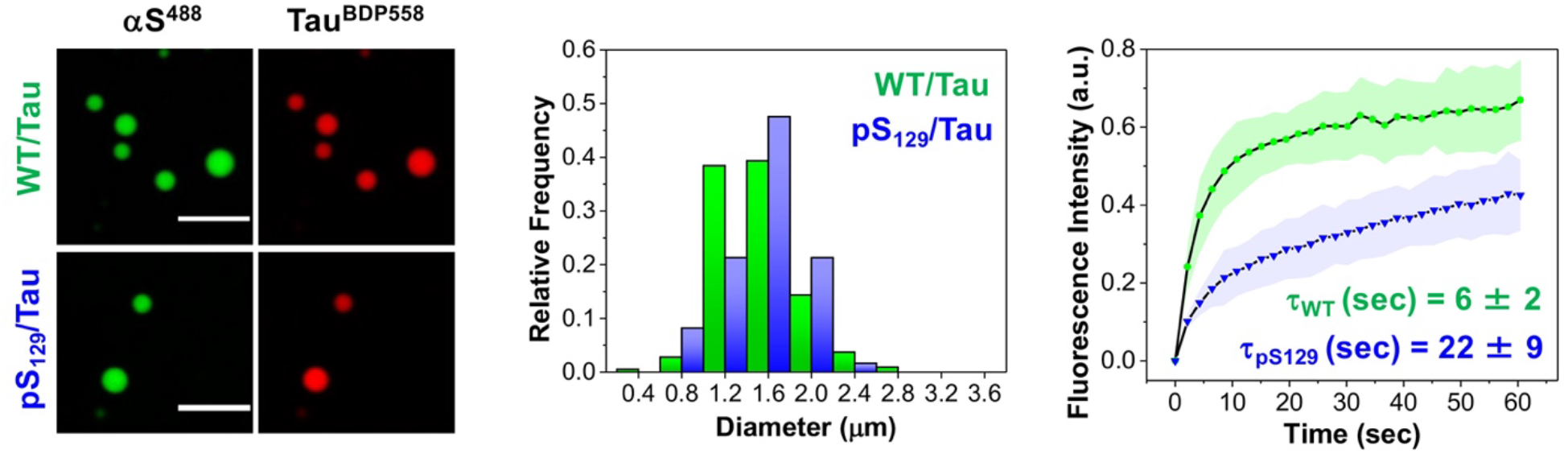
Effect of pS_129_ on size distribution and dynamics of αS in αS/tau co-condensates. Left: Representative images of WT/tau and pS_129_/tau (40 μM αS /20 μM tau) condensates in LLPS buffer, doped with 5% αS-C^488^_9_ or 5% αS-C^488^_9_pS_129_, and 5% tau^BDP558^. Middle: Size distribution of WT/tau and pS_129_/tau after 1 h of incubation. Condensate diameters were measured using Image J. Right: FRAP profile showing the translational mobility of WT or pS_129_ in αS/tau co-condensates (doped with 5% αS-C^488^_9_ or 5% αS-C^488^_9_pS_129_). Error bars represent the standard deviation of 3 measurements. Ex: 488 nm and Em: 500-540 nm for Alexafluor 488-labeled αS, Ex: 561 nm and Em: 570-620 nm for tau^BDP558^. All experiments were performed in an LLPS buffer: 20 mM HEPES, 150 mM NaCl, and 1 mM TCEP at pH 7.4, with 15% PEG-8000, at room temperature. Scale bars are 5 μm.

## Conclusion

In this work, we have described the production and characterization of αS and tau with authentic PTMs introduced by GCE or enzymatic modification. These methods allow for site-specific incorporation of phosphorylation and acetylation, while being broadly accessible to biochemistry laboratories that may lack peptide synthesis expertise. Additionally, we demonstrated the utility of these approaches for isotopic and fluorescent labeling, allowing for structural and biophysical studies. While Phos-tag gel electrophoresis provided a useful tool for assessing phosphorylation status, we also highlighted the need for careful interpretation, since band shifts can be influenced by the intrinsic charge of the phosphorylated region as observed for αS-pS_129_. Beyond protein production, we examined the biophysical and functional consequences of these modifications, particularly focusing on phosphorylation at Ser87, Ser129, and Tyr39 in αS and acetylation at Lys274, Lys280, or Lys281 in tau.

By integrating radioligand binding studies with PTM-modified fibrils, we demonstrated how PET probe interactions can serve as a functional readout of structural changes induced by modifications. Our findings suggest that pY_39_, when present at physiological sub-stoichiometric ratios, perturbs PET probe binding only at Site 2, likely reflecting blocked interactions by phosphorylation and localized, but not global, structural changes of the fibrils. Future work involving detailed structural characterization will be essential for understanding the underlying molecular basis of this altered binding affinity. Additionally, since pY_39_ interacts with K_21_ and K_34_ in the 6l1t/6l1u structure, investigating the impact of Lys acetylation at these sites on fibril conformation and radioligand binding will also be of interest, particularly if both PTMs can be incorporated into the same construct.

We also examined how PTMs influence αS’s native functions, including membrane interactions. Vesicle-binding studies using NMR and FCS showed that pS_87_ does not significantly impact αS-vesicle interactions. This finding contrasts with previous studies using phosphomimetic mutations or enzyme-blocking mutations, which showed that this PTM reduces helicity of the protein on vesicles (38). This highlights the importance of employing authentic modifications rather than relying on mimics when studying PTM effects. As noted above, investigation of Lys acetylation in αS is also of interest, and this charge-neutralizing PTM would be expected to inhibit αS interactions with vesicles, at least for some sites. In the case of tau acetylation, we observed a greater impairment in polymerization rate for ^Ac^K_280_/^Ac^K_281_ compared to ^Ac^K_274_, consistent with literature reports suggesting that the ^Ac^K_280_ and ^Ac^K_281_ sites might have a more pathological role. One interpretation of acetylation effects is that they prevent ubiquitination and thus protein degradation while also impairing interaction with microtubules, making tau more accessible for phosphorylation and subsequent aggregation. Indeed, Yan et al. found that inhibition of the deacetylase HDAC6 by tubastatin A increased tau KXGS motif phosphorylation at S262 (50). Acetylation at KXGS tau motifs may inhibit phosphorylation, suggesting a protective role of one PTM over another, potentially leading to aggregation (29; 45). Investigation of such cross-talk between PTMs is another exciting opportunity afforded by our GCE methods, and efforts are underway to apply the pSer-3.1G system to tau phosphorylation.

Finally, we demonstrated that pS_129_ modification does not affect the size of αS/tau co-condensates but significantly reduces αS mobility, showing a role in modulating condensate dynamics. Additionally, the impact of pS_129_ modification on αS dynamics provides a foundation for understanding how phosphorylation influences transitions from condensates to mature fibrils, highlighting a role in the underlying mechanisms of amyloid aggregation.

Overall, this work showcases the value, as well as some challenges, of GCE and enzymatic methods for producing PTM-modified proteins and their application in biophysical and structural characterization. With these tools, we provide new insights into the role of acetylation and phosphorylation in αS and tau, contributing to a deeper understanding of PTM-mediated regulation in neurodegenerative disease pathology. There are also clear opportunities to use GCE and enzymatic modification together to achieve incorporation of two or more PTMs in the same protein, which will allow for exploration of the most impactful and physiologically-relevant combinations of PTMs. Finally, while we have focused on avoiding methods like NCL to ensure that these techniques are as accessible as possible, using GCE and enzymatic modifications in combination with chemical peptide synthesis and NCL can allow one to efficiently access very highly modified proteins, as we have shown in some contexts with αS and look forward to systematic applications in both αS and tau (47; 23).

## AUTHOR CONTRIBUTIONS

Ibrahim G. Saleh: Investigation (lead); writing – original draft (lead); writing – review and editing (lead). Marie Shimogawa: Conceptualization (lead); investigation (lead); writing – original draft (supporting). Jennifer Ramirez: Conceptualization (lead); investigation (lead); writing – original draft (lead). Bernard Abakah: Investigation (supporting). Yarra Venkatesh: Investigation (supporting); writing – review and editing (supporting); resources (supporting). Honey Priya James: Investigation (supporting); resources (supporting). Ming-Hao Li: Investigation (supporting); resources (supporting). Sarah A. Louie: Investigation (supporting). Marshall G. Lougee: Investigation (supporting). Wai-Kit Chia: Investigation (supporting); resources (supporting). Richard Cooley: Supervision (supporting); writing – review and editing (supporting). Ryan A. Mehl: Funding acquisition (supporting); supervision (supporting); writing – review and editing (supporting). Tobias Baumgart: Funding acquisition (supporting); supervision (supporting); writing – review and editing (supporting). Robert Mach: Funding acquisition (supporting); supervision (supporting); writing – review and editing (supporting). David Eliezer: Funding acquisition (supporting); supervision (supporting); writing – review and editing (supporting). Elizabeth Rhoades: Conceptualization (lead); supervision (lead); funding acquisition (lead); writing – review and editing (lead). E. James Petersson: Conceptualization (lead); supervision (lead); funding acquisition (lead); writing – original draft (lead); writing – review and editing (lead).

## ACKNOWLEDGMENTS

This research was funded by the University of Pennsylvania and by the National Institutes of Health (NIH RF1-NS125770 to David Eliezer, Elizabeth Rhoades, and E. James Petersson supporting αS PTM studies; NIH RF1-NS103873 to E. James Petersson supporting αS/tau interaction studies, NIH U19-NS110456 to E. James Petersson and Robert H. Mach supporting αS PET tracer development, NIH R01-GM097552 to Tobias Baumgart supporting membrane studies, and NIH RM1-GM144227 to the GCE4All Biomedical Technology Development and Dissemination Center at Oregon State University). Marie Shimogawa thanks the Nakajima Foundation for scholarship funding. Jennifer Ramirez was supported by the NIH Chemistry Biology Interface Training Program (NIH T32-GM133398). Marshall G. Lougee was supported by an Age Related Neurodegenerative Disease Training Grant fellowship (NIH T32-AG000255). Instruments supported by the NIH and NSF include NMR (NSF CHE-1827457) and mass spectrometers (NIH S10-OD030460).

## DATA AVAILABILITY STATEMENT

The data that support the findings of this study are available from the corresponding author upon reasonable request.

