## Supporting Information for "Genetic Code Expansion and Enzymatic Modifications as Accessible Methods for Studying Site-Specific Post-Translational Modifications of Alpha-Synuclein and Tau"

**Contents**

List of Figures, Schemes, Tables, and Equation....S3
General Information ....S4
Construction of αS Expression Plasmids ....S6
Constructs of Tau Expression Plasmids ....S6
αS pSer Protein Production ....S8
αS $pY_{39}$ Protein Production ....S9
Production of αS-$pS_{129}$ and αS-$C_9pS_{129}$ by PLK2 Kinase Co-Expression ....S10
Fluorescent Labeling of αS ....S11
Tau Protein Production ....S12
Fluorescent Labeling of Tau ....S13
Production of $^{Ac}$K Tau ....S13
Radioligand Saturation Binding ....S15
Preparation of Synthetic Vesicles ....S16
Heteronuclear Single Quantum Coherence Spectroscope (HSQC) ....S16
Fluorescence Correlation Spectroscopy (FCS) ....S17
Calculation of Vesicle Binding Affinity ....S18
Tubulin Purification ....S20
Tubulin Polymerization Assays ....S20
αS/tau Condensate Imaging ....S21
Fluorescence Recovery After Photobleaching (FRAP) ....S21
References ....S35

**Figures:**

Figure S1. SDS-PAGE purification and Phos-tag gel GCE αS-p$S_{87}$ purification .... S23
Figure S2. HPLC chromatograms GCE αS-p$S_{87}$ .... S24
Figure S3. MALDI-MS GCE αS-p$S_{87}$ .... S25
Figure S4. SDS-PAGE gel and MALDI-MS αS-p$S_{87}$ fluorescent labeling .... S26
Figure S5. MALDI-MS and MS/MS αS c-Abl p$Tyr_{39}$ trypsin digest .... S27
Figure S6. Gel, HPLC, and MALDI-MS αS-$C_9$ and enzymatic coexpression αS-C9p$S_{129}$ .... S28
Figure S7. Gel, FPLC, and MALDI-MS GCE $Tau_{0N4R}$ $^{Ac}K_{274}$ and $^{Ac}K_{280}$ .... S29
Figure S8. Gel and MALDI-MS $Tau_{0N4R}$-WT fluorescent labeling .... S30
Figure S9. HSQC spectra GCE αS-p$S_{87}$ .... S31
Figure S10. Representative FCS curves GCE αS-p$S_{87}C_{114}^{Atto488}$ with and without 275µM lipid .... S32
Figure S11. Representative images of condensate FRAP .... S33
Figure S12. Representative images of wide-field condensate droplet size .... S34

**Tables:**

Table S1: DNA Sequences for cloning .... S6

**Equations:**

Equation S1. Global fitting for total binding and nonspecific binding of radioligand .... S15
Equation S2. FCS One-component fit .... S18
Equation S3. FCS Two-component fit .... S18
Equation S4. Vesicle binding curve .... S19
Equation S5. FRAP ROI .... S22
Equation S6. FRAP ratio of bleached to unbleached regions .... S22
Equation S7. FRAP recovery profile single exponential model .... S22

### General Information

*E. coli* BL21(DE3) cells and *E. coli* Dh5α cells were purchased from New England Biotechnologies (Ipswich, MA, USA). DNA oligomers were purchased from Integrated DNA Technologies, Inc (Coralville, IA, USA). DNA extraction and Miniprep kits were purchased from Qiagen (Hilden, Germany). Buffers were made with MilliQ filtered (18 MΩ) water (Millipore; Billerica, MA, USA). Preparation of the pTXB1-αS-intein-$H_6$ plasmid containing α-synuclein (αS) with a C-terminal fusion to the *Mycobacterium xenopi* GyrA intein and C-terminal $His_6$ tag was described previously (1). This plasmid was used as a starting point for the preparation of αS constructs. The 0N4R tau sequence was cloned into a lab-made pET-HT vector (a gift from L. Regan). All tau sequences contained an N-terminal $His_6$ tag with a TEV protease cleavage site. This was the starting point for the preparation of all tau constructs.

For acetyllysine incorporation, the pTECH-chAcK3RS (IPYE) plasmid was a gift from David Liu via Addgene (plasmid # 104069; http://n2t.net/addgene:104069; RRID:Addgene_104069; Watertown, MA, USA). Acetyllysine was purchased from ChemImpex. Nicotinamide was purchased from Alfa Aeser (Tewksbury, MA, USA).

For phosphoserine incorporation, we used the pKW2-EFSep plasmid, developed by Ryan Mehl and Rick Cooley (Plasmid #173897, https://www.addgene.org/173897/) (14). An *E. coli* bacterial strain with an RF1 and SerB deletion named B95(DE3) ΔA ΔfabR ΔserB (bacterial strain #197655, https://www.addgene.org/197655/) was used for the phosphoserine genetic code expansion system. For $^{15}N$ isotopic labeling of protein, ammonium chloride ($^{15}N$, 99% pure) and Celtone base powder ($^{15}N$, 98% pure) were purchased from Cambridge Isotope Labs (Tewksbury, MA, USA). For Phos-tag gels, the Phos-tag Acrylamide was purchased from Avantor Science Central (Radnor, PA).

Atto488-maleimide was purchased from Sigma-Aldrich. Alexa Fluor 488 was purchased from ThermoFisher. BODIPY 558/568 maleimide (BDP558-maleimide) was purchased from LumiProbe.

Matrix-assisted laser desorption/ionization mass spectrometer (MALDI-MS) data were collected with a Bruker RapifleX MALDI-MS instrument or a Bruker Microflex MALDI-MS (Billerica, MA, USA). UV/Vis absorbance spectra were obtained with a Hewlett-Packard 8452A diode array spectrophotometer (currently Agilent Technologies; Santa Clara, CA). Gel images were obtained with a Syngene G:Box mini-6 (Cambridge, UK). Proteins were purified on a 1260 Infinity II preparative high-performance liquid chromatography (HPLC) system (Agilent Technologies). Proteins were also purified on a Cytiva ÄKTA pure FPLC system using a HiLoad Superdex 200 pg size exclusion column (Marlborough, MA, USA).

For radioligand binding assays, a Unifilter-96 harvesting system (Perkin Elmer; Waltham, MA, USA) was used with MicroScint-20 scintillation cocktail (PerkinElmer) and counted on the Microbeta system (Perkin Elmer).

For vesicle binding assays, Large Unilamellar Vesicles (LUVs) were prepared with a 50:50 ratio of 1-palmitoyl-2-oleoyl-glycero-3-phosphocholine (POPC) and 1-palmitoyl-2-oleoyl-sn-glycero-3-phospho-L-serine (POPS), both purchased from Avanti Research (Alabaster, AL). The vesicles were then extruded using the LiposoFast-Basic & Stabilizer model purchased from Avestin, Inc. (Ottowa, ON, Canada). The 50 nm pore membranes were purchased from Cytiva (Marlborough, MA, USA). DLS was collected on the ZetaSizer Nano Series Nano-ZS purchased from Malvern Panalytical (Westborough, MA).

For protein condensate imaging and FRAP experiments, confocal microscopy was performed using an Olympus IX83 inverted microscope equipped with a FluoView 3000 scanning system (Olympus, Center Valley, PA). Imaging was conducted at room temperature using a 60× 1.2 NA water-immersion objective lens (Olympus). Image analysis was carried out using ImageJ software (version 1.52a). For imaging droplets in solution, glass coverslips ($25 \times 25$ $mm^2$) were obtained from Fisher Scientific (Waltham, MA).

### Construction of αS Expression Plasmids

The following primers (**Table S1**) were designed for site-directed mutagenesis to introduce TAG (= Z) codons at each Lys acetylation or Ser phosphorylation site. Site-directed mutagenesis for αS TAG mutations were performed on the pTXB1-αS-intein-$H_6$ plasmid described previously (1). The parent αS plasmid contains αS fused to a His-tagged GyrA intein from *Mycobacterium xenopi* (αS-intein) and was cloned into a pRBC vector (provided by OSU) using Gibson Assembly with the primers given in **Table S1** to avoid compatibility issues with their origins of replication.

### Construction of Tau Expression Plasmids

Site-directed mutagenesis for the tau TAG mutations were performed on the pET-$H_6$-TEV-Tau0N4R plasmid. The parent tau plasmid contains tau fused to a $His_6$ tag with a TEV protease cleavage site before the tau0N4R sequence. To produce acetylated tau constructs, plasmids encoding tau0N4R with a TAG (= Z) mutation at the target site were generated. Primers used for cloning are listed in **Table S1**. Numbering of tau0N4R mutations is based on full-length tau2N4R protein [UniProt:P10636-8]).

**Table S1.** DNA Sequences for cloning:

| | | |
|---|---|---|
| αS-$Z_{87}$ | Forward | 5’-GGGAGCAGGGTAGATTGCAGCAG -3’ |
| | Reverse | 5’-TCCACTGTCTTCTGGGCT -3’ |
| αS-$Z_{129}$ | Forward | 5’-GAAATGCCTTAGGAGGAAGGGTATC-3’ |
| | Reverse | 5’-ATAAGCCTCATTGTCAGG-3’ |
| αS-$C_{114}$ | Forward | 5’-GGAATTCTGTGCGATATGCCT-3’ |
| | Reverse | 5’-GGAATTCTGTGCGATATGCCT-3’ |
| Tau-$Z_{274}$ | Forward | 5’-GGAGGCGGGTAGGTGCAGATAAT-3’ |
| | Reverse | 5’-GGGCTGGTGCTTCAGGTTCT-3’ |

| | | |
|---|---|---|
| Tau-$Z_{280}$ | Forward | 5’-ATAATTAACTAGAAGCTGGATCTTAGC-3’ |
| | Reverse | 5’-CTGCACCTTCCCGCCTCC-3’ |
| Tau-$Z_{281}$ | Forward | 5’-GATAATTAATAAGTAGCTGGATCTTAG-3’ |
| | Reverse | 5’-TGCACCTTCCCGCCTCCC-3’ |
| Gibson | Forward | 5’- GCTGCTAAACAAAGCCCGAAAGGAAGCTGAGTTGGCT-3’ |
| | Reverse | 5’-ATACATCCATATGTATATCTCCTTCTTAAAGTTAAACAAAATTATTT CTAGAG-3’ |

G-block
AGATATACAtATGGATGTATTCATGAAAGGACTTTCAAAGGCCAAGGAGGGAGTTGTGGCTG
CTGCTGAGAAAACCAAACAGGGTGTGGCAGAAGCAGCAGGAAAGACAAAAGAGGGTGTTC
TCTATGTAGGCTCCAAAACCAAGGAGGGAGTGGTGCATGGTGTGGCAACAGTGGCTGAGAA
GACCAAAGAGCAAGTGACAAATGTTGGAGGAGCAGTGGTGACGGGTGTGACAGCAGTAGCC
CAGAAGACAGTGGAGGGAGCAGGGAGCATTGCAGCAGCCACTGGCTTTGTCAAAAAGGACC
AGTTGGGCAAGAATGAAGAAGGAGCCCCACAGGAAGGAATTCTGGAAGATATGCCTGTGGA
TCCTGACAATGAGGCTTATGAAATGCCTTCTGAGGAAGGGTATCAAGACTATGAACCTGAAG
CCTGCATCACGGGAGATGCACTAGTTGCCCTACCCGAGGGCGAGTCGGTACGCATCGCCGAC
ATCGTGCCGGGTGCGCGGCCCAACAGTGACAACGCCATCGACCTGAAAGTCCTTGACCGGC
ATGGCAATCCCGTGCTCGCCGACCGGCTGTTCCACTCCGGCGAGCATCCGGTGTACACGGTG
CGTACGGTCGAAGGTCTGCGTGTGACGGGCACCGCGAACCACCCGTTGTTGTGTTTGGTCGA
CGTCGCCGGGGTGCCGACCCTGCTGTGGAAGCTGATCGACGAAATCAAGCCGGGCGATTAC
GCGGTGATTCAACGCAGCGCATTCAGCGTCGACTGTGCAGGTTTTGCCCGCGGGAAACCCGA
ATTTGCGCCCACAACCTACACAGTCGGCGTCCCTGGACTGGTGCGTTTCTTGGAAGCACACC
ACCGAGACCCGGACGCCCAAGCTATCGCCGACGAGCTGACCGACGGGCGGTTCTACTACGC
GAAAGTCGCCAGTGTCACCGACGCCGGCGTGCAGCCGGTGTATAGCCTTCGTGTCGACACGG
CAGACCACGCGTTTATCACGAACGGGTTCGTCAGCCACGCTACTGGCCTCACCGGTCTGAAG
CTTCATCATCATCATCATCATTAATGAgGTACCCTTGGGACAGAGTATCCCCCGCAGGCGGA
AGTAACGCTCTGGTGCCACGCGGTAGTAAAGAAACCGCTGCTGCTAAATTCGAACGCCAGC
ACATGGACAGCTCTACTTCTGCTGCTCTCGAGGCTTAATTAACCTAGGCTGCTAAACAAAGC
CCGAA

**αS pSer Protein Production**

Expression of αS with pSer incorporated at residue S87 or S129 followed the method previously described (13). The plasmid encoding αS with a TAG mutation at the site of interest (backbone: pRBC, origin of replication: p15A, antibiotic resistance: Amp) and a machinery plasmid for phosphoserine incorporation, pkW2-EFSep (origin of replication: pBR322, antibiotic resistance: Chlor) were co-transformed by heat shock at 42 °C into BL21 (DE3) competent *E. coli* cells with a Δ*serB* genomic knockout. Cells were plated and incubated on Amp/Chlor plates. Single colonies were picked to inoculate primary cultures in 50 mL ZY-non inducing media (NIM) supplemented with 0.1 mg/mL Amp and 0.025 mg/mL Chlor, in a 250 mL baffled flask. Primary cultures were incubated overnight at 37 °C with 250 rpm shaking.

To generate non-isotopically labeled, Ser-phosphorylated αS, secondary cultures in ZY-auto-inducing media (AIM) ($OD_{600}$ ~ 5-8) were inoculated by adding 1% inoculum of ZY-NIM culture and grown at 37 °C with shaking at 250 rpm until the $OD_{600}$ reached ~ 1.5. The culture was subsequently cooled to 22 °C and grown with shaking at 250 rpm.

To generate isotopically labeled αS-$pS_{87}$, protein expression followed the method previously described (12). Secondary cultures were first started in fresh ZY-NIM media with 10% inoculum and grown at 37 °C with shaking at 250 rpm until $OD_{600}$ reached ~3-4. Cells were then pelleted by centrifugation at 4000 g for 10 min. Cell pellets were resuspended in an equal volume of minimal MIM 2 media that contains $^{15}$N-ammonium chloride and $^{15}$N-CELTONE. We note that for this work, we did not supplement L-serine. This was grown at 37 °C with shaking at 250 rpm in a baffled flask until the $OD_{600}$ increased by 1-2 units. Expression of the gene of interest was induced with 1 mM isopropyl ß-D-1-thiogalactopyranoside (IPTG). Induced cells were then grown in the shaker-incubator at 22 °C overnight.

In all the buffers used during affinity purification, phosphate inhibitors (20 mM sodium fluoride, 5 mM sodium pyrophosphate, 0.5 mM sodium orthovanadate, final concentration) were added. After centrifugation (4000 rpm, 30 min, 4 °C), cell pellets were re-suspended in buffer (40 mM Tris pH 8.3, 1 Roche protease inhibitor tablet, 1 mM phenylmethylsulfonyl fluoride) and sonicated in a cup in an ice-water bath (5 min, 1 s ON, 2 s OFF). The resulting lysate was centrifuged (14,000 rpm, 30 min, 4 °C), and supernatant containing the protein of interest (POI) was purified over a Ni-NTA affinity column. Intein cleavage was carried out by incubation with 200 mM βME on a rotisserie overnight at room temperature. Cleaved protein was dialyzed into 20 mM Tris, pH 8 buffer before purification over a second Ni-NTA column to remove the free intein from the sample. The αS proteins were purified by RP-HPLC over a C4 column, dialyzed into 1x phosphate buffered saline (PBS) and spin-concentrated. For purification of phosphorylated Cys mutants for fluorescent labeling, TCEP was added to a final concentration of 1 mM prior to HPLC. Upon flash-freezing, protein stocks were kept at -80 °C until further use. Isotopically labeled αS samples were lyophilized after HPLC purification. Phos-tag gels were run to analyze %phosphorylation. The Phos-tag gels were prepared with a top 4% acrylamide top stacking layer, and a 12% acrylamide layer. Phos-tag was added to the 12% layer at a 33.3 μM concentration, along with 66.7 μM $MnCl_2$.

**αS $pY_{39}$ Protein Production**

The procedures for purifying the catalytic domain of c-Abl Tyr kinase as well as phosphorylation of αS were adapted from our previously published work (10).

To express and purify c-Abl for phosphorylation studies, BL21 (DE3) *E. coli* cells were transformed with pET $His_6$-SUMO-TEV-cAbl (Amp resistance) and YopH (Strep resistance) plasmids. Colonies were selected on LB agar plates containing ampicillin and streptomycin, and primary cultures were grown overnight. Secondary cultures were inoculated and grown in LB medium until reaching an $OD_{600}$ of 0.6–0.7, at which point they were cooled to 16 °C, induced with 0.25 mM IPTG, and incubated overnight. Cells were harvested by centrifugation and lysed by sonication in 50 mM Tris, 500 mM NaCl, 5% glycerol, pH 8.0, supplemented with protease inhibitors. The lysate was cleared by centrifugation at

14,000 rpm for 30 min, and the supernatant was incubated with a Ni-NTA column equilibrated in 50 mM HEPES pH 7.5, supplemented with 5% glycerol. After sequential washes, c-Abl was eluted using an imidazole gradient (60–300 mM). The eluted fractions were analyzed by SDS-PAGE, pooled, and incubated with TEV protease for SUMO tag cleavage during overnight dialysis into 20 mM Tris pH 8.0, supplemented with 5% glycerol and 1 mM TCEP. Following cleavage confirmation, the sample was run on anion exchange chromatography with a 0–350 mM sodium chloride gradient. To remove TEV and further purify c-Abl, the sample was run through size-exclusion chromatography (SEC) using an S75 or S200 column, followed by concentration to 50–150 μM. Aliquots were flash-frozen and stored at -80 °C until further use. Co-expression of YopH was essential to counteract c-Abl toxicity in *E. coli,* and a gradient Ni-NTA purification step was employed to improve separation of c-Abl from YopH.

To generate αS-pY$_{39}$, wild type (WT) αS was prepared in 50 mM Tris, 150 mM NaCl, pH 7.5, at a final concentration of 50–100 μM. c-Abl was added at a molar ratio of 0.0436 equivalents relative to αS, following an established literature protocol (4). The reaction was initiated by the addition of Mg-ATP (100 mM) and $MgCl_2$ (1 M) and incubated at 30 °C for 2–4 hours. Phosphorylation progress was monitored by MALDI-MS, and the reaction was halted when approximately 50% modification was achieved. Occasionally, degraded αS was observed, likely due to non-specific cleavage by contaminating TEV. To remove unreacted c-Abl and separate phosphorylated from unmodified αS, the reaction mixture was concentrated and subjected to either SEC (Superdex Increase 75 column) or HPLC (C4 column). The eluted fractions were analyzed by MALDI-MS to confirm phosphorylation and ensure product purity. The final products were characterized using phos-tag SDS-PAGE and analytical HPLC.

Additionally, a c-Abl plasmid where TEV recognition sequence is removed so that SUMO could be cleaved more specifically with a SUMO protease Ulp-1 was created. Further investigations are necessary to evaluate the utility of this strategy.

**Production of αS-p$S_{129}$ and αS-$C_9$p$S_{129}$ by PLK2 Kinase Co-expression**

αS-p$S_{129}$ was generated and purified as described above with the inclusion of a kanamycin-resistant plasmid encoding for polo-like kinase 2 (PLK2) (gift from D.T.S. Pak), a kinase that targets serine 129 (11). A 1.0 L LB broth was supplemented with 100 μg/mL ampicillin and 34 μg/mL chloramphenicol and induced with 1 mM IPTG at OD ~0.6 overnight at 16 °C. The culture was centrifuged at 4600x g for 20 minutes at 4 °C. The pellet was resuspended in 30 mL of 40 mM Tris pH 8 supplemented with 2 tablet of EDTA-free protease inhibitor tablet (Roche), and 0.1 mM PMSF. The resuspended pellet was then sonicated on ice for 2 minutes, 1 second on/2 seconds off, power set to 50 W. The cellular debris was removed by centrifugation at 20,000 x g for 30 minutes. The supernatant (after centrifugation) was filtered with a 0.22 μM syringe filter and added to 5–7 mL of Ni-NTA resin equilibrated with αS-intein buffer 1 (ASB1; 50 mM HEPES pH 7.5), followed by incubation with rocking at 4 °C for ~1 hour. The resin was then washed with ~15 mL ASB1, followed by a wash with αS-intein buffer 2 (ASB2; 50 mM HEPES pH 7.5, 5 mM imidazole). The protein was eluted with ~12 mL Ni-NTA αS-intein buffer 3 (ASB3: 50 mM HEPES pH 8, 400 mM imidazole). To cleave the His-tagged intein, β-mercaptoethanol (BME; 200 mM final concentration) was added to the eluate and incubated overnight (16–18 hours) at room temperature with rocking. The sample was dialyzed against 20 mM Tris pH 8.0 at 4 °C with three changes of the dialysis buffer. Removal of the cleaved His-tagged intein, as well as any un-cleaved αS-intein, was achieved by incubation of the cleaved sample with Ni-NTA resin equilibrated with 20 mM Tris pH 8.0 for 1 hour at 4 °C with rocking. The column flow through (containing cleaved αS) was filtered using a 0.22 μm filter and further purified using HPLC over a C4 preparatory column. Phosphorylation was confirmed via mass shift with MALDI-TOF using a Bruker rapifleX. αS-$C_9$p$S_{129}$ was generated and purified in an identical fashion. We made an $S_9$C mutation to attach Alexa Fluor 488 (Af488) via maleimide chemistry

**Fluorescent Labeling of αS**

To fluorescently label αS Cys mutants, the protein stocks in 20 mM Tris, 50 mM NaCl, pH 7.4 were incubated with 2-10 eq. TCEP, then 10 eq. Atto488-maleimide dye was added and the reaction tube was wrapped with aluminum foil and incubated at room temperature for 2-4 hours and further incubated at 4 °C overnight with stirring until product formation was observed by MALDI-MS. The labeled protein was exchanged into 20 mM Tris pH 7.4, 50 mM NaCl buffer, and unreacted dye was removed by passing the solution over two coupled HiTrap Desalting columns. Following purification, the sample concentration was determined using a NanoDrop™ spectrophotometer using the following extinction coefficients: Alexa 488 ε@494 nm = 73000 $M^{-1}cm^{-1}$, αS ε@280 nm = 5960 $M^{-1}cm^{-1}$ and calculated by correcting the absorbance signal at 280 nm by Alexa 488 using the following equation: A(αS)280 = [A280-0.11(A494)]/A280). The samples were then aliquoted into microcentrifuge tubes, flash frozen, and stored at -80 °C.

**Tau Protein Production**

A 1 L tau expression for 0N4R tau in LB broth (Miller) supplemented with 100 µg/mL ampicillin was induced with 1 mM IPTG at OD ~0.6 for 4 hours at 37 °C. The culture was then pelleted at 4600 rpm for 20 minutes at 4 °C. The pellet was resuspended in 30 mL of Ni-NTA tau Buffer A (TBA: 50 mM Tris pH 8, 500 mM NaCl, 10 mM imidazole) with 1 mg/mL chicken egg-white lysozyme (Sigma), 1 tablet of EDTA-free protease inhibitor tablet (Roche), and 1 mM phenylmethylsulfonyl fluoride (PMSF). The resuspended pellet was sonicated on ice for 1 minute 40 seconds, 1 second on/2 seconds off, power set to 50 W. The cellular debris was removed by centrifugation at 20,000 x g for 30 minutes. The supernatant (after centrifugation) was filtered with a 0.22 µm syringe filter and added to 5-7 mL of Ni-NTA resin equilibrated with TBA, followed by incubation with rocking at 4 °C for ~1 hour. The resin was then washed with ~30 mL TBA and the protein was eluted with ~15 mL Ni-NTA tau Buffer B (TBB: 50 mM Tris pH 8, 500 mM NaCl, 400 mM imidazole). The eluent was then exchanged back into TBA and concentrated to ~1 mL using Amicon concentrators (Sigma). The His-tag was cleaved by overnight incubation at 4 °C with TEV protease (100 µL of 260 µM added to 1 mL tau) and freshly prepared 1 mM dithiothreitol (DTT).

Removal of the cleaved His-tag, as well as any uncleaved His-tagged tau, was achieved by incubation of the cleaved sample with TBA equilibrated Ni-NTA resin for 1 hour at 4 °C with rocking. The column flow-through containing the cleaved tau protein was exchanged into tau Buffer C (TBC; 25 mM Tris PH 8, 100 mM NaCl) using an Amicon concentrator and concentrated down to 0.5-2 mL. The solution was filtered using a 0.22 μm syringe filter and further purified on an ÄKTA pure FPLC system using a HiLoad Superdex 200 pg size exclusion column. The protein was aliquoted out into microcentrifuge tubes, flash frozen, and stored at -80 °C.

**Fluorescent Labeling of Tau**

For generating labeled constructs, freshly purified tau was reduced by incubation with 1 mM DTT for 10 minutes. DTT was removed by exchanging the sample into labeling buffer (20 mM Tris pH 7.4, 50 mM NaCl, 6 M guanidine HCl) using Amicon filters. BDP558-maleimide in DMSO was added in five-fold molar excess to protein and incubated overnight at 4 °C with stirring. Labeled protein was exchanged into 20 mM Tris pH 7.4, 50 mM NaCl buffer, and unreacted dye was removed by passing the solution over two coupled HiTrap Desalting columns. Following purification, the samples concentration was determined using a NanoDrop™ spectrophotometer using the following extinction coefficients: BDP558 ε@561 nm = 84400 $M^{-1}cm^{-1}$, $tau_{0N4R}$ ε@280 nm = 7450 $M^{-1}cm^{-}1$, and calculated by correcting the absorbance signal at 280 nm by BDP558 using the following equation: $A(tau)_{280}= [A_{280}-0.11(A_{561})]/A_{280})$, and then were aliquoted out into microcentrifuge tubes, flash frozen and stored at -80 °C.

**Production of $^{Ac}$K Tau**

Tau-$Z_{274}$, Tau-$Z_{280}$, or Tau-$Z_{281}$ plasmids and the machinery plasmid for acetyl-lysine incorporation, pTECH-chAcK3RS (IPYE), were co-transformed into BL21 (DE3) competent cells via heat shock at 42 °C for 45 seconds. These cells were then plated on ampicillin/chloramphenicol (Amp/Chlor) agar plates and incubated. Single colonies were selected to inoculate primary cultures in LB media supplemented with 0.1 mg/mL Amp and 0.025 mg/mL Chlor, and these cultures were grown overnight or until turbid at 37 °C. Secondary cultures in LB media were inoculated and grown at 37 °C with shaking at 250 rpm until the optical density reached approximately 0.6. The culture was then kept at 37 °C, followed by the addition of 50 mM nicotinamide and 10 mM ε-acetyl lysine. After a 5-minute incubation, the expression of the target gene was induced with 1 mM isopropyl β-D-1-thiogalactopyranoside (IPTG). The constructs were then purified as described above for WT tau. Acetylation was confirmed via MALDI-TOF.

**Radioligand Saturation Binding**

αS fibril samples were prepared by shaking 100 μM αS (total protein concentration) at 1500 rpm for 72 hours. Fibrils were resuspended in buffer then diluted to 50 nM and incubated for 1 h at 37 °C with increasing concentrations of [$^3$H]Tg-190b or [$^3$H]BF-2846. Total binding is measured with the absence of competitors and non-specific binding is defined by the presence of 5 uM unlabelled Tg-190b and 300 nM unlabeled BF-2846, respectively. The fibrils were incubated with ten different concentrations of [$^3$H]Tg-190b (1.56-fold dilution, 0.7 nM – 40 nM) and [$^3$H]BF-2846 (1.56-fold dilution, 0.5 nM – 20 nM). The mixture with a total reaction volume of 150 μL is incubated in 50 mM Tris-HCl buffer with 0.01% BSA for 1.5 hours at 37 °C in a non-binding 96 well plate (Corning). After incubation, bound and free radioligand is harvested with Unifilter-96 harvesting system (Perkin Elmer), followed by washing with an ice-cold buffer containing 10 mM Tris-HCl (pH 7.4), 150 mM NaCl and 20 % EtOH. Filters containing bound ligands are added with 50 μL scintillation cocktail (MicroScint-20, PerkinElmer) and counted on the Microbeta system (Perkin Elmer). All data points were acquired in triplicate. The equilibrium dissociation constant ($K_d$) and the maximal number of binding sites ($B_{max}$) were determined by globally fitting the total binding and nonspecific binding data to Equation S1:

$$Y = B_{max} * \left(\frac{X}{K_d + X}\right) + NS * X \qquad \text{(S1)}$$

where X is the concentration of tritiated ligand and NS is nonspecific binding, using GraphPad Prism software (San Diego, CA, USA).

**Preparation of Synthetic Vesicles**

Lipid vesicles were prepared by extrusion through porous membranes. A mixture in 50:50 molar ratio of 1-palmitoyl-2-oleoyl-sn-glycero-3-phosphoserine (POPS) and 1-palmitoyl-2-oleoyl-sn-glycero-3-phosphocholine (POPC) were drawn from chloroform stock and dried under nitrogen gas to form a film inside a glass vial. Films were desiccated under vacuum and re-hydrated in 20 mM 3-(*N*-morpholino) propanesulfonic acid (MOPS), 147 mM NaCl, 2.7 mM KCl, pH 7.4. Ten freeze-thaw cycles consisting of cooling in liquid nitrogen for 40 s and warming in a 60 °C water bath for 2 min were performed to aid the formation of uniformly sized vesicles. With syringes, vesicles were then extruded 31 times through stacked 50 nm pore membranes held in place inside an extruder. Vesicles were determined by dynamic light scattering (DLS) to be monodisperse and distributed uniformly around 80 nm in diameter, consistent across different concentrations of all samples. All lipid vesicles were prepared fresh and used within 48 h of extrusion.

**Heteronuclear Single Quantum Voherence Spectroscopy (HSQC)**

Lyophilized WT or lysine-acetylated αS mutants were dissolved in NMR buffer (10 mM $Na_2HPO_4$, 100 mM NaCl, 10% v/v $D_2O$, pH 6.8) and mixed in a 1:1 volume ratio with SUV stock solution or NMR buffer. Final protein concentrations were ca. 100 µM and final SUV lipid concentrations were ca. 3 mM, assuming complete conversion of lipids to SUVs. NMR $^1H$-$^{15}N$ heteronuclear single quantum coherence (HSQC) experiments were acquired at 10 °C using a 500 MHz Bruker Avance spectrometer equipped with a cryogenic probe using TopSpin 3.2 software. Data processing and analysis were performed using NMRbox (7), NMRPipe (3), and NMRFAM-SPARKY (6) software. Amide resonance peaks were assigned based on previously published chemical shift assignments for WT αS monomer (5). NMR peak intensity ratios were calculated as the ratio of peak intensity in the presence of SUVs to that in the absence of SUVs for matched samples. To correct for protein concentration variations in the samples, the intensity ratios were normalized by the average ratio for the C-terminal 40 residues, which do not feature appreciable membrane interactions at these lipid concentrations.

**Fluorescence Correlation Spectroscopy (FCS)**

Fluorescence correlation spectroscopy (FCS) experiments to study the binding of $\alpha$S-$C^{488}{}_{114}$ and $\alpha$S-$pS_{87}C^{488}{}_{114}$, including preparation of synthetic lipid vesicles, collection of FCS data, and data analysis, were carried out as described previously for arginylated αS (9). Eight-well chambered coverglasses (Nunc, Rochester, NY, USA) were prepared by plasma cleaning followed by incubation overnight with polylysine-conjugated polyethylene glycol (PEG-PLL), prepared using a modified Pierce PEGylation protocol (Pierce, Rockford, IL, USA). PEG-PLL coated chambers were rinsed with and stored in Milli-Q water until use. FCS measurements were performed on a lab-built instrument based on an Olympus IX71 microscope with a continuous emission 488 nm DPSS 50 mW laser (Spectra-Physics; Santa Clara, CA, USA). All measurements were made at 20 °C. The laser power entering the microscope was adjusted to 4.5 µW. Fluorescence emission collected through the objective was separated from the excitation signal through a Z488rdc long pass dichroic filter and an HQ600/200m bandpass filter (Chroma; Bellows Falls, VT, USA). Emission signal was focused onto the aperture of a 50 µm optical fiber. Signal was amplified by an avalanche photodiode (Perkin Elmer; Waltham, MA, USA) coupled to the fiber. A digital autocorrelator (Flex03Q-12, correlator.com; Bridgewater, NJ, USA) was used to collect 10 autocorrelation curves of 10 seconds for each measurement of free protein in buffer without lipids or 30 autocorrelation curves of 30 seconds for each measurement in the presence of lipid vesicles. Fitting was done using lab-written code in MATLAB (The MathWorks; Natick, MA, USA).

To determine the diffusion time of each protein construct, each αS variant labeled with Atto 488 was measured in buffer without lipid. The average of 10 autocorrelation curves was fit to a 1-component autocorrelation function:

$$G(\tau) = \frac{1}{N}\left(\frac{1}{1+\frac{\tau}{\tau_1}} * \left(\frac{1}{1+\frac{s^2\tau}{\tau_1}}\right)^{\frac{1}{2}}\right) \quad \text{(S2)}$$

where $G(\tau)$ is the autocorrelation function, N is the number of molecules in the focal volume, $\tau_1$ is the diffusion time of αS, and s is the radial-to-axial ratio of the excitation volume. The counts per molecule (CPM) for each sample was calculated by dividing the average intensity (Hz) of the measured signal by the number of molecules N. The normalized CPM of each αS was calculated by dividing by the CPM of freely diffusing fluorescent standard Alexa Fluor 488.

**Calculation of Vesicle Binding Affinity**

αS constructs labeled with Atto488 were examined in the presence of varying concentrations (0.001 mM to 0.5 mM lipid) of lipid vesicles consisting of 50:50 POPS/POPC. The average of 30 autocorrelation curves was fit to a 2-component equation:

$$G(\tau) = \frac{1}{N}\left(A * \frac{1}{1+\frac{\tau}{\tau_1}} * \left(\frac{1}{1+\frac{s^2\tau}{\tau_1}}\right)^{\frac{1}{2}} + Q * (1-A) * \frac{1}{1+\frac{\tau}{\tau_2}} * \left(\frac{1}{1+\frac{s^2\tau}{\tau_2}}\right)^{\frac{1}{2}}\right) \quad \text{(S3)}$$

where $G(\tau)$ is the autocorrelation function, N is the number of molecules in the focal volume, $\tau_1$ is the characteristic diffusion time of αS, $\tau_2$ is the characteristic diffusion time of the vesicles, s is the radial-to-axial ratio of the excitation volume, Q is the ratio of the brightness of vesicle-bound αS relative to αS, and A is the fraction of free αS. When fitting the autocorrelation curves for αS in the presence of lipid vesicles, the diffusion time of bound and unbound αS were respectively fixed to experimentally determined values. The diffusion time of unbound protein, $\tau_1$, was determined by measurements of the protein in buffer without lipids. Since bound protein diffuses with the vesicles to which they are bound, the diffusion time of the vesicles, $\tau_2$, was determined by measurements of the protein in the presence of a concentration of vesicles that gave the maximum diffusion time (0.1 mM lipid). In the binding assay, the fraction of αS bound at each lipid concentration was obtained from the fit to each autocorrelation curve. Averages and standard deviations were calculated from at least 3 independent measurements performed on separate days at each lipid concentration. The resulting binding curve was fit to the following equation, from which the $K_{d,app}$ was determined.

$$A = \frac{B_{max}X}{K_{d,app}+X} \qquad \text{(S4)}$$

Where A is the fraction of αS bound, x is the accessible lipid concentration, $B_{max}$ is the maximum fraction of αS bound, and $K_{d,app}$ is the apparent dissociation constant.

**Tubulin Purification**

Tubulin was purified from young bovine brains as previously described (2), with two cycles of polymerization and depolymerization, with one modification: the concentration of nucleotides was adjusted to 1.5 mM ATP and 1 mM GTP for the first polymerization, then increased to 2.5 mM ATP and 1.5 mM GTP for the second polymerization step. The purified tubulin was flash-frozen in liquid nitrogen in BRB80 buffer (80 mM PIPES, 1 mM $MgCl_2$, 1 mM EGTA, pH 6.8) and stored at -80 °C. When needed, aliquots were quickly thawed at room temperature, then clarified by centrifugation at 100,000xg for 6 minutes at 4 °C. Tubulin concentration was determined by measuring the absorbance at 280 nm with a molar extinction coefficient of 115,000 $M^{-1}cm^{-1}$, and the clarified tubulin was used within 2 hours.

**Tubulin Polymerization Assays**

The polymerization of soluble tubulin into microtubules was monitored by measuring the increase in scattered light at 340 nm, as previously described (8), with modifications for microplate adaptation. Tubulin aliquots were clarified by centrifugation and left in BRB80. For each reaction, 20 μM tubulin was incubated with 10 μM tau for 2.5 minutes on ice. After the addition of 1 mM GTP (Sigma), the final reaction consists of BRB80 buffer supplemented with fresh 1 mM DTT, 20 μM tubulin, 10 μM tau, 1 mM GTP at 200 μL in microcentrifuge tubes on ice. Samples were then immediately transferred to a pre-warmed 96 well optical Thermo Sci Nunc plate (full area, black plate, clear bottom) and measured for 60 minutes at 37 °C in a microplate reader (M1000, Tecan) with both excitation and emission at 340 nm. The samples were then returned to 4 °C for 5 minutes to confirm the absence of protein aggregation through cold depolymerization. These measurements were performed in triplicate and averaged to obtain statistical variations. Polymerization curves were normalized via min-max re-scaling (0 to 1) for WT and acetylated tau constructs, while GTP/tubulin only samples were normalized to the max of WT.

**αS/tau Condensate Imaging**

The desired protein condensate solution (25 μL) was prepared using either WT αS or $pS_{129}$-modified αS (40 μM), containing 5% αS-$C_9^{Af488}$ (WT) or 5% αS-$C_9^{Af488}pS_{129}$ ($pS_{129}$). This was mixed with $Tau_{0N4R}$ (20 μM), which included 5% $tau^{BDP558}$, in a liquid-liquid phase separation (LLPS) buffer consisting of 20 mM HEPES, 150 mM NaCl, and 1 mM TCEP, adjusted to pH 7.4. A 15% (w/v) PEG 8000 solution was then added. The reaction mixture was gently mixed by pipetting up and down to ensure homogeneity, followed by incubation at room temperature for 30 minutes before imaging.

Confocal microscopy imaging was performed using an Olympus IX83 inverted microscope, equipped with FluoView3000 scanning system (Olympus, Center Valley, PA). Images were taken at room temperature using a 60x 1.2 NA water immersion objective lens (Olympus). Imaging of multi-protein systems was performed *via* orthogonal fluorescence labeling of proteins using Alexafluo 488 ($\lambda_{ex}$ 488 nm, $\lambda_{em}$ 500–540 nm) and BDP558 ($\lambda_{ex}$ 561 nm, $\lambda_{em}$ 570–620 nm). The excitation lasers were alternated to minimize crosstalk between the different channels using a sequential line scan mode. Images were analyzed with ImageJ (version 1.52a). For imaging droplets in solution, glass coverslips (25 × 25 $mm^2$, Fisher Scientific) were passivated with BSA (2 mg/mL in HEPES buffer). Solutions (10 μL) containing the droplets incubated for 30 min and were imaged in a closed chamber created by sandwiching two coverslips (25 × 25$mm^2$, Fisher Scientific) using vacuum grease.

**Fluorescence Recovery After Photobleaching (FRAP)**

FRAP experiments of WT/tau or $pS_{129}$/tau (40 μM/20 μM) condensates were performed with the same confocal set up as described above. A circular region of interest (ROI) within a protein condensate settled on the glass coverslip was bleached using short exposures (~60 s) of 488 nm lasers at 50% laser power. The collection of images for the recovery stage started immediately after the bleaching. FRAP data were analyzed using ImageJ (for image intensity extraction) and Microsoft Excel (for quantitative analysis). For each time frame, mean intensities were estimated for an ROI within the bleached region and of another ROI from the unbleached reference region. The ratio of intensities at bleached ($I_{bleach}(t)$) and reference regions ($I_{ref}(t)$) was determined at each time point and normalized to 1 for the intensity ratio before photobleaching ($q(t_{prebleach})$) and to 0 for the intensity ratio at 0 s after photobleaching ($q(t_0)$) using the following formula:

$$I_{norm}(t) = \frac{q(t) - q(t_0)}{q(t_{prebleach}) - q(t_0)} \quad \text{(S5)}$$

$$\text{where } q(t) = \frac{I_{bleach}(t)}{I_{ref}(t)} \quad \text{(S6)}$$

The normalized intensities were plotted against time to obtain a fluorescence recovery profile. The recovery profile was fit to a single exponential model to obtain the fluorescence recovery rate, $\tau_{FR}$.

$$I(t) = A\left(1 - e^{-\frac{t}{\tau_{FR}}}\right) \quad \text{(S7)}$$

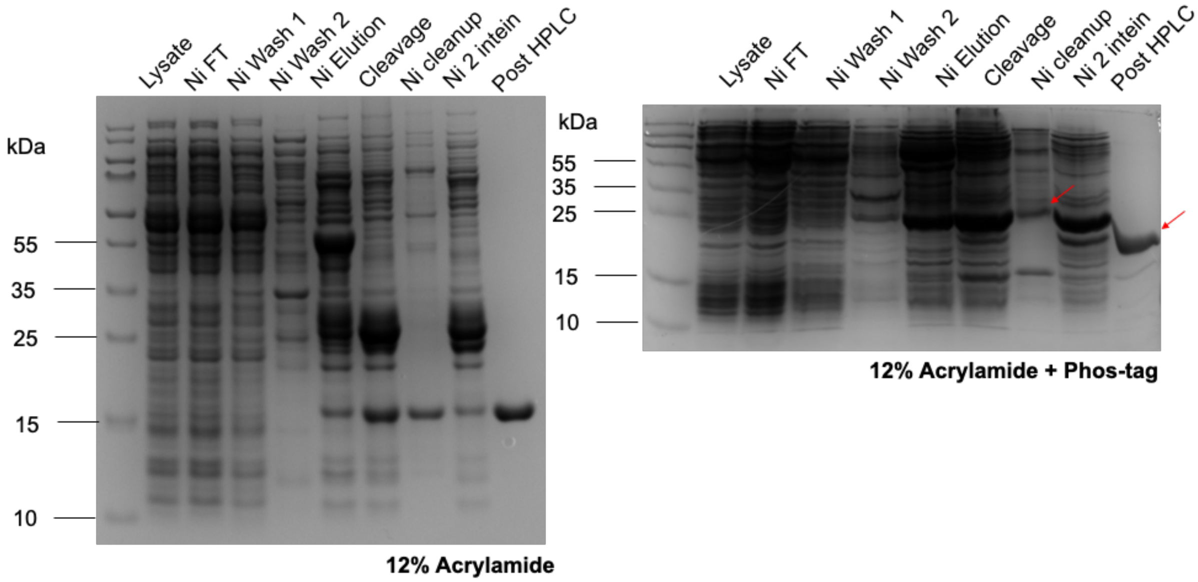


**Figure S1**. Affinity purification of αS with phosphorylation at S87, produced by GCE. SDS-PAGE gels show affinity purification of αS-pS$_{87}$ from BL21 ΔserB cells, both run with 12% acrylamide and stained with Coomassie blue. Left = 12% acrylamide, Right = 12% acrylamide and Phos-tag.

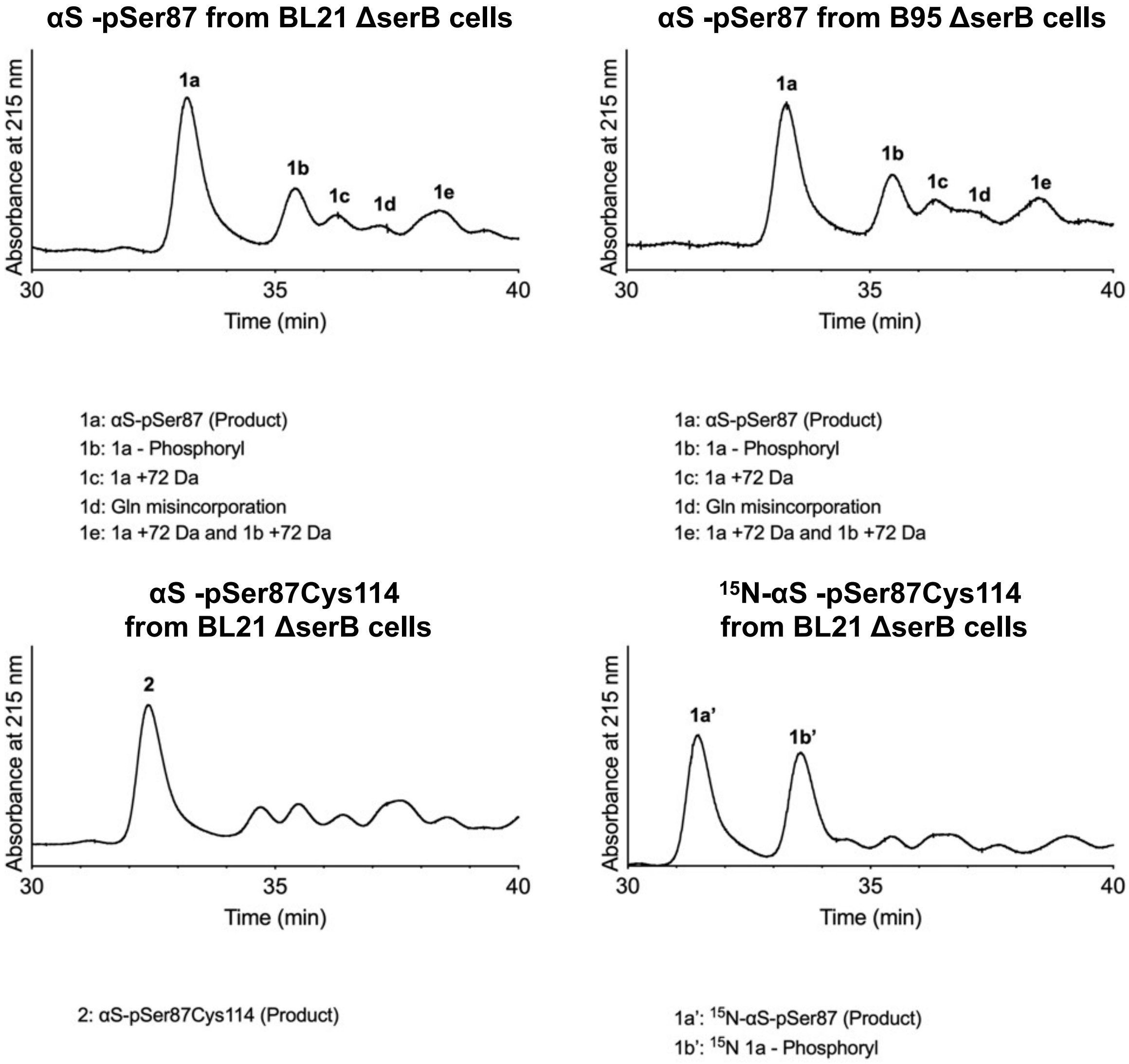


**Figure S2.** HPLC chromatogram collected for affinity-purified phosphoserine GCE products. Affinity-purified αS-p$S_{87}$ from BL21 ΔserB cells or B95 ΔserB cells, αS-p$S_{87}C_{114}$ and $^{15}$N αS-p$S_{87}$ from BL21 ΔserB cells. Peak identity was confirmed by MALDI-MS.

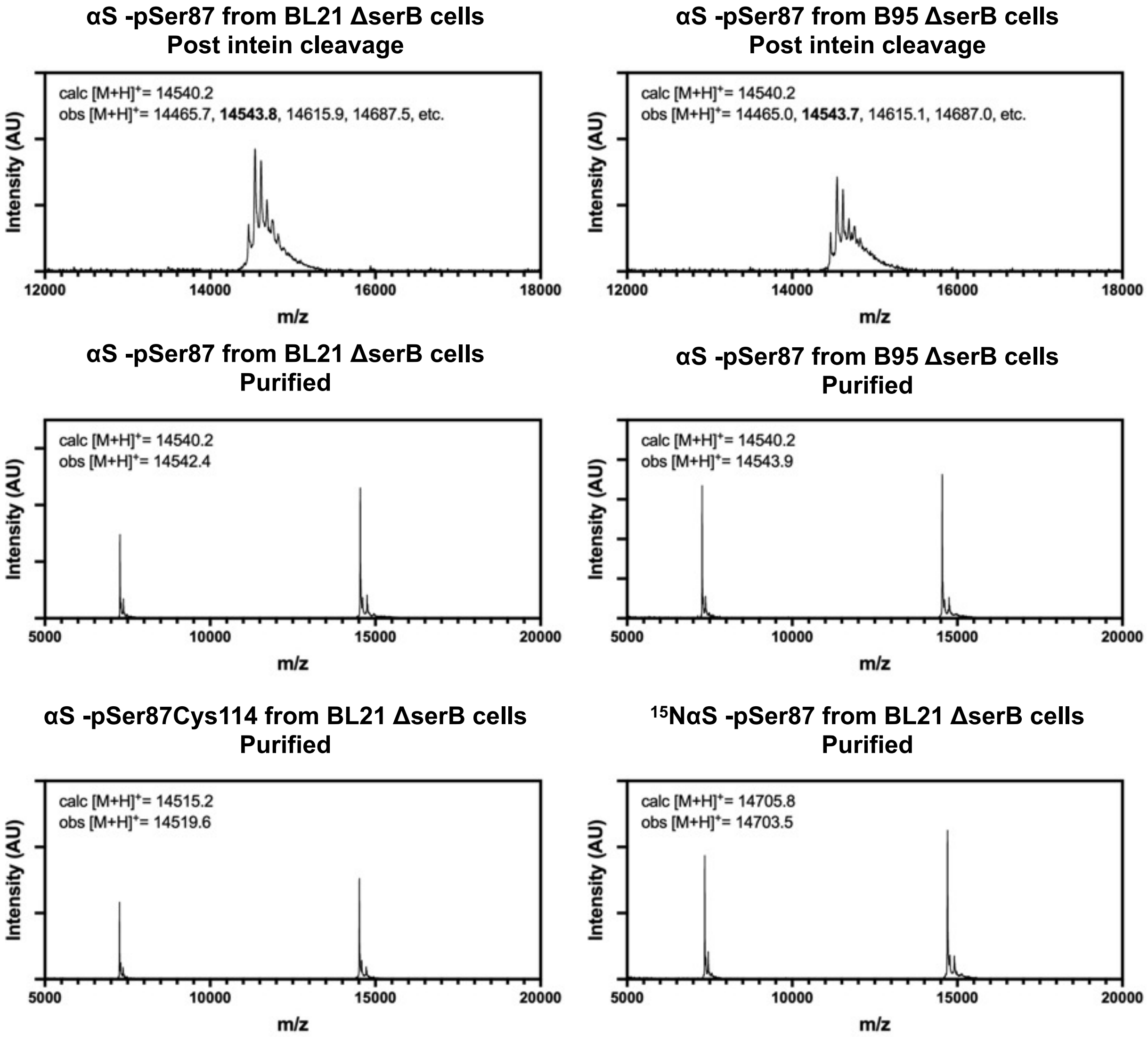


**Figure S3.** MALDI-MS characterization of affinity-purified or HPLC-purified αS-pS87 constructs.

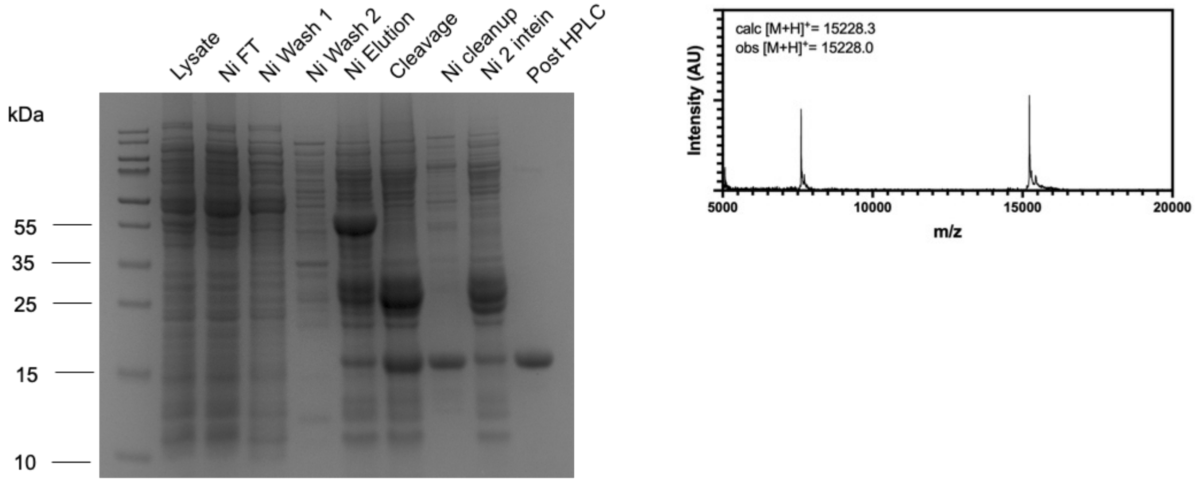


**Figure S4.** Preparation of fluorescently labeled αS-p$S_{87}$. Left: SDS-PAGE analysis of affinity purification and HPLC purification of αS-p$S_{87}C_{114}$ Right: MALDI-MS characterization of HPLC-purified αS-p$S_{87}C_{114}^{Atto488}$.

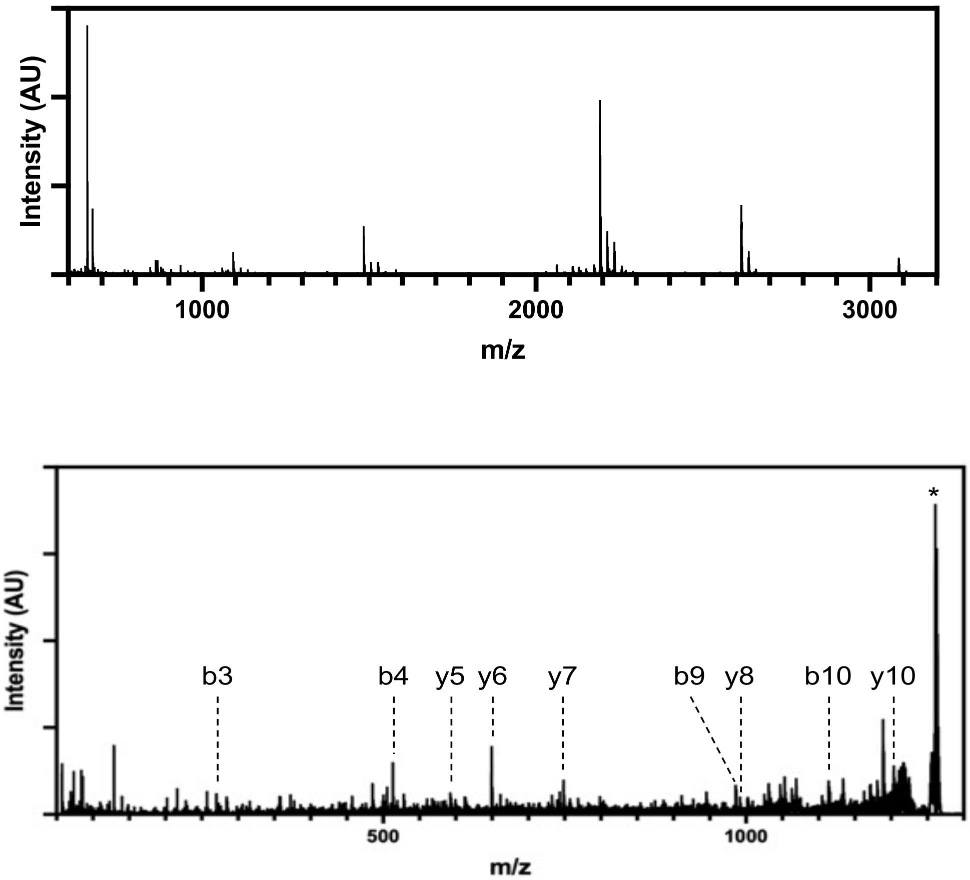


| b | | | | y |
|---|---|---|---|---|
| --- | 1 | G | 11 | --- |
| 157.0972 | 2 | V | 10 | 1203.6021 |
| 270.1812 | 3 | L | 9 | 1104.5337 |
| 513.2109 | 4 | Y(Phospho) | 8 | 991.4496 |
| 612.2793 | 5 | V | 7 | 748.4199 |
| 669.3008 | 6 | G | 6 | 649.3515 |
| 756.3328 | 7 | S | 5 | 592.3301 |
| 884.4277 | 8 | K | 4 | 505.2980 |
| 985.4754 | 9 | T | 3 | 377.2031 |
| 1113.5704 | 10 | K | 2 | 276.1554 |
| --- | 11 | E | 1 | 148.0604 |

**Figure S5.** Analysis of αS phosphorylated by Tyr kinase c-Abl. Top: Trypsin digest of αS after 2 h kinase treatment. Bottom: MALDI-MSMS data for the αS $_{36\text{-}46}$ peptide.

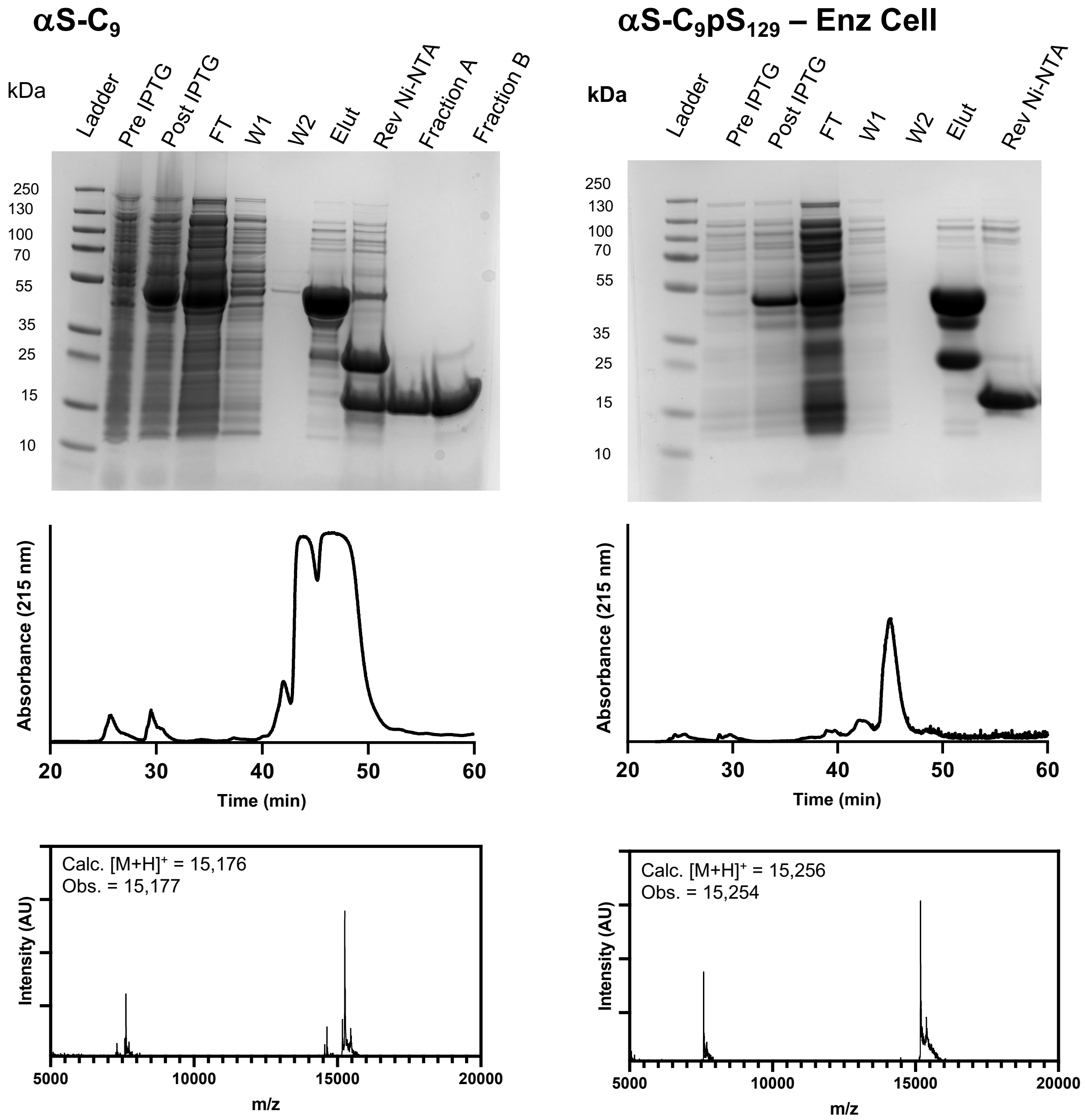


**Figure S6.** Gel, HPLC, and MALDI-MS analysis of αS-$C_9$ and αS-$C_9$p$S_{129}$ from enzymatic co-expression.

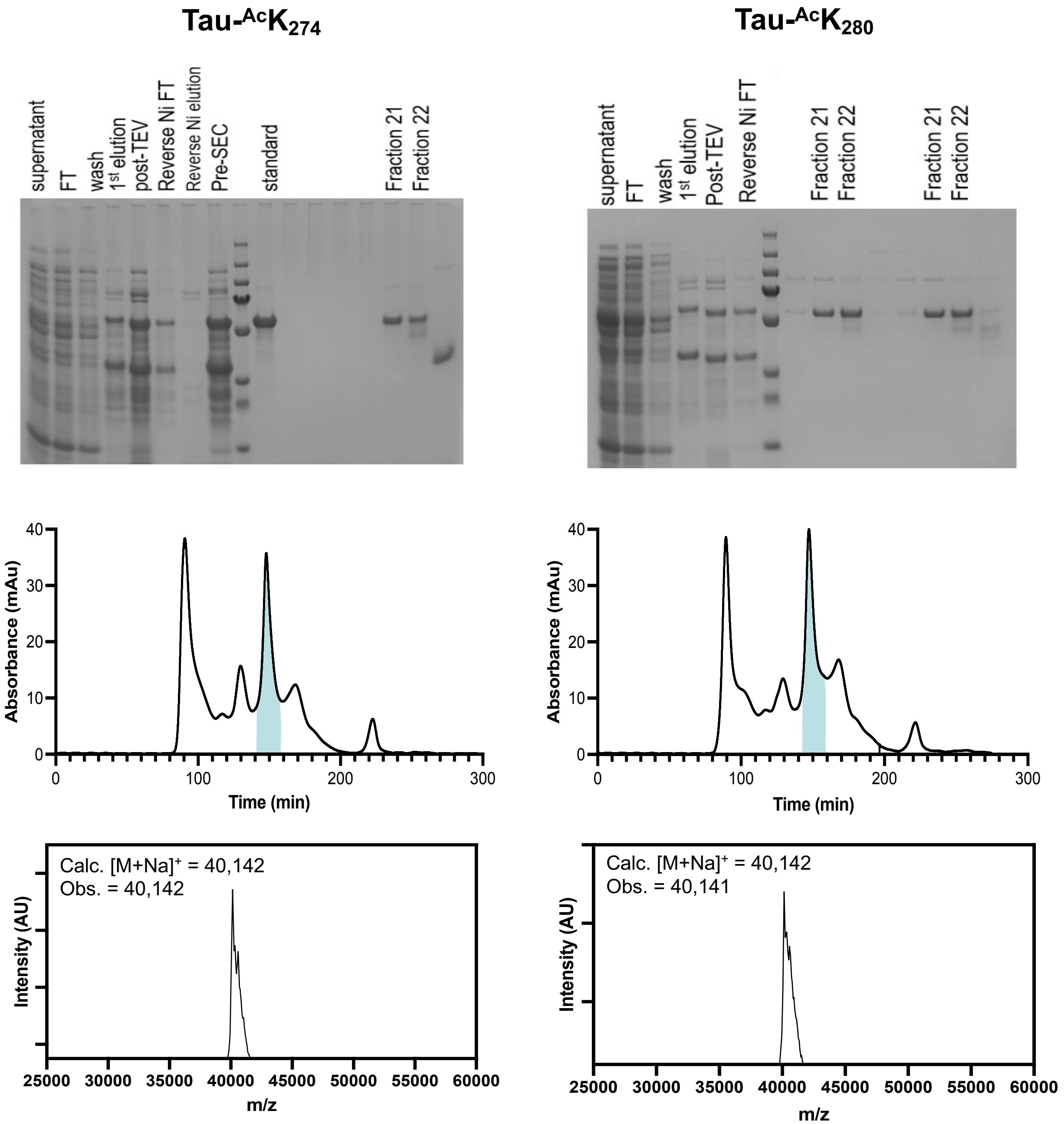


**Figure S7.** Gel, FPLC, and MALDI-MS analysis of 0N4R tau $^{Ac}K_{274}$, and $^{Ac}K_{280}$.

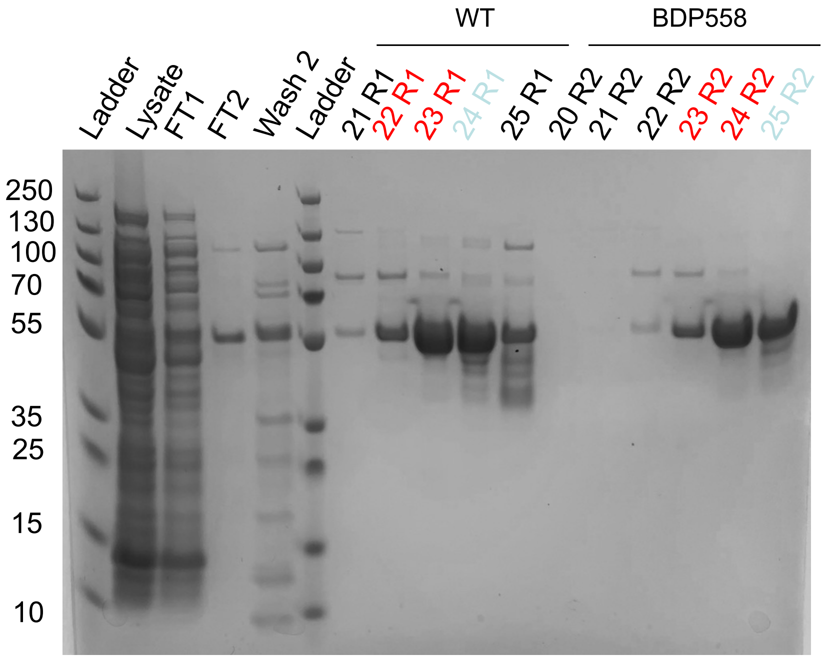


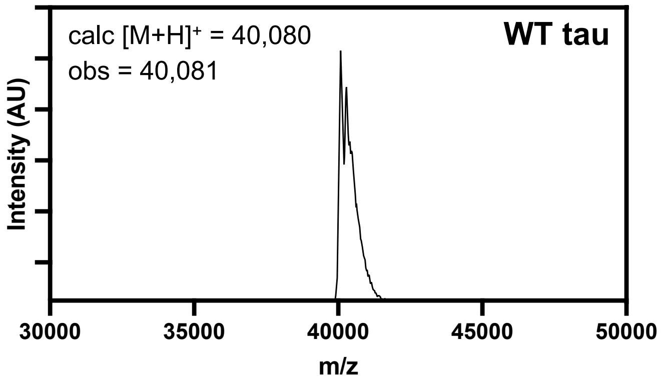


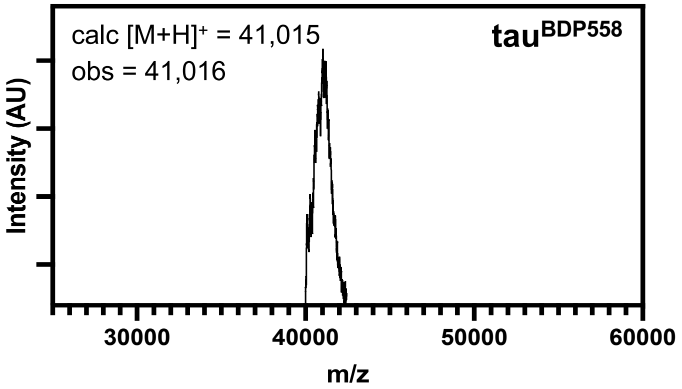


**Figure S8.** Gel and MALDI-MS analysis of labeled tau construct.

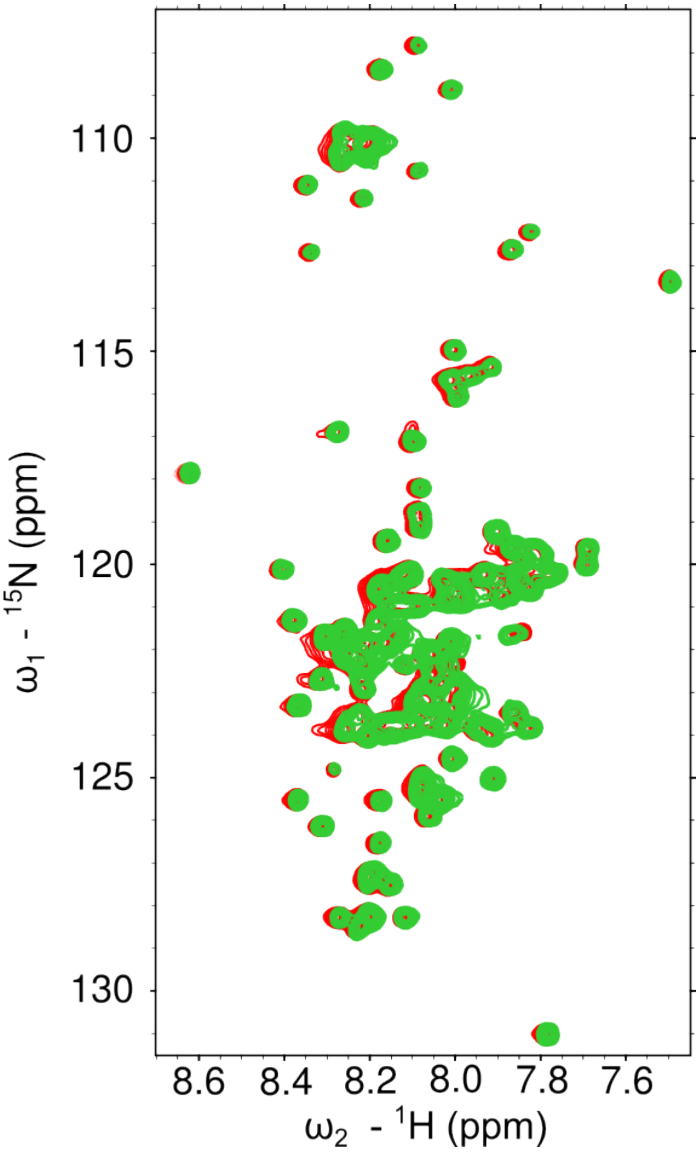


**Figure. S9**. $pS_{87}$ HSQC spectra with (red) and without (green) vesicles.

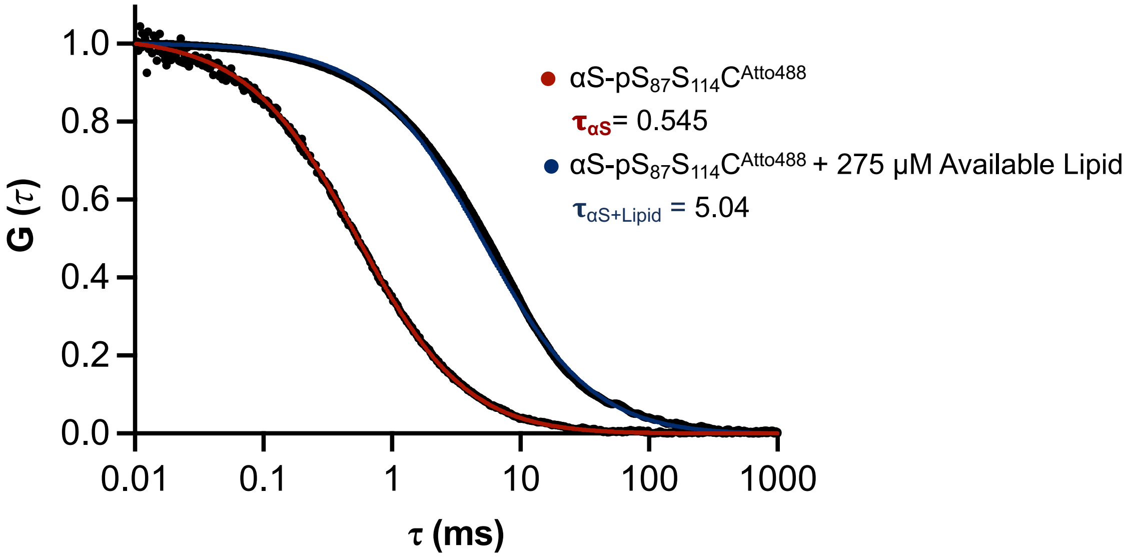


**Figure S10.** Representative normalized averaged data and autocorrelation curves of aS-$pS_{87}C_{114}^{Atto488}$ (n = 10) and aS-$pS_{87}C_{114}^{Atto488}$ +275 μM Available Lipid (n = 30).

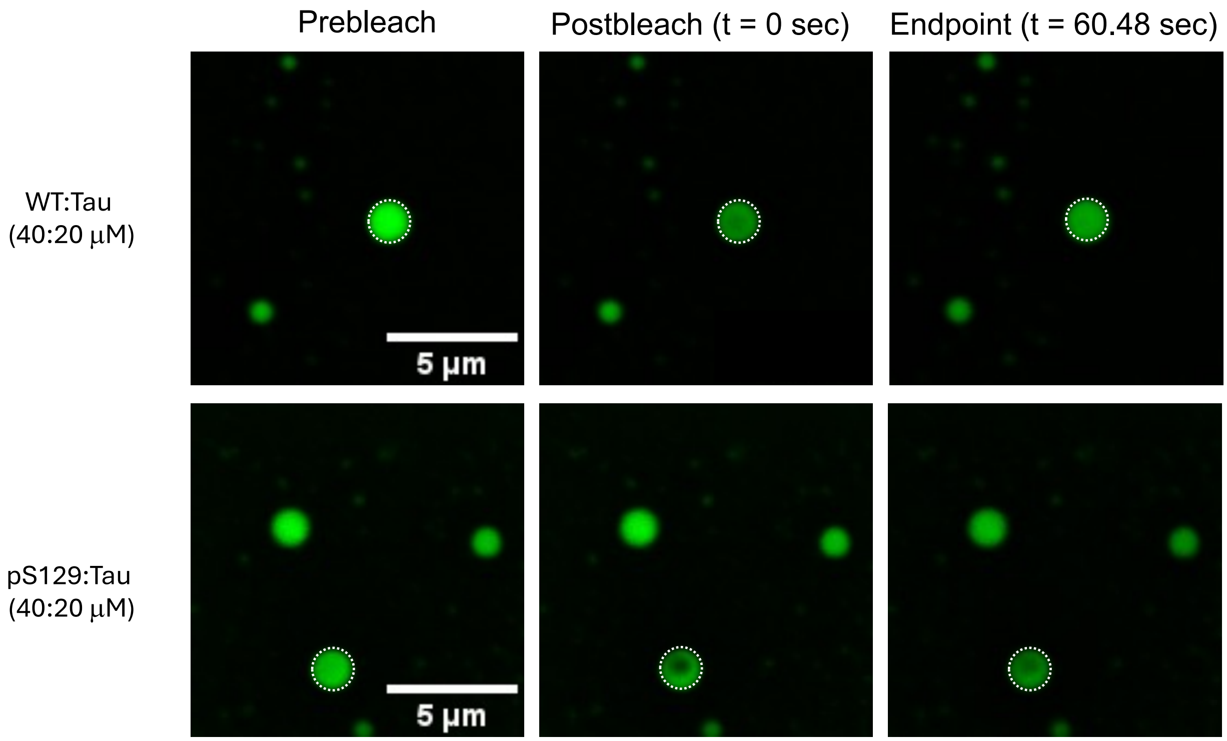


**Figure S11**. Representative images of fluorescence recovery after photobleaching (FRAP) of αS within WT/tau (top) and $pS_{129}$/tau (bottom) condensates doped with 5% αS-$C_9^{Af488}$ or 5% αS-$C_9^{Af488}pS_{129}$. Ex: 488 nm and Em: 500-540 nm for αS-$C_9^{Af488}$ or αS-$C_9^{Af488}pS_{129}$. The incubation time for WT/tau condensates is 30 min before imaging.

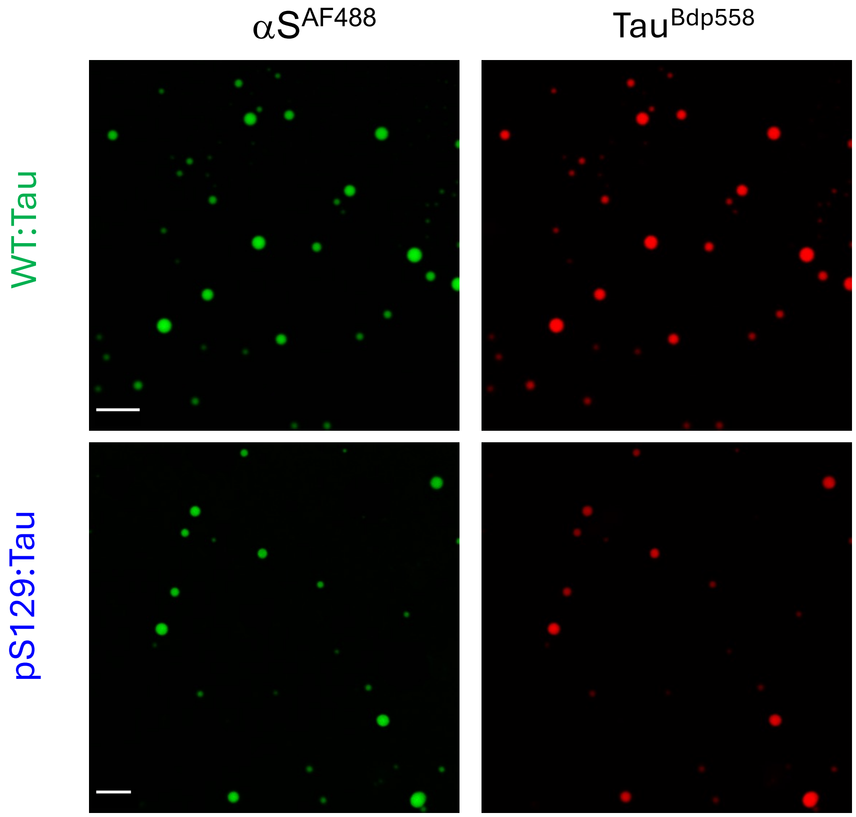


**Figure S12**. Representative images showing wide fields of view used in quantifying droplet size distribution for αS within WT/tau (top) and $pS_{129}$/tau (bottom) condensates doped with 5% αS-$C_9^{Af488}$ or 5% αS-$C_9^{Af488}$$pS_{129}$. Ex: 488 nm and Em: 500-540 nm for αS-$C_9^{Af488}$ or αS-$C_9^{Af488}$$pS_{129}$. The incubation time for WT/tau condensates is 30 min before imaging. Scale bar is 5 µm.
